# CGK733 binds to adenine nucleotide translocator 2 and modulates mitochondrial function and protein translation

**DOI:** 10.1101/2025.11.13.688169

**Authors:** Daisuke Kaida, Yuta Inagaki, Keisuke Yaku, Kazuki Sasaki, Sakurako Kuwata Ikeda, Seiichi Koike, Masami Shima, Shintato Iwasaki, Tomonao Inobe, Takashi Nakagawa, Minoru Yoshida, Ken Ishigami, Ryo Katsuta

**Author notes:** Correspondence: Daisuke Kaida Faculty of Medicine, Academic Assembly, University of Toyama, 2630 Sugitani, Toyama 930-0194, Japan.

## Abstract

Chemical genetics is a powerful strategy for dissecting biological mechanisms, and a crucial step in this approach is to identify the molecular target responsible for a compound’s activity. CGK733, a compound with anti-proliferative activity, was once reported as an ATM/ATR inhibitor, although this activity has been debated, and its target and mode of action have remained unclear. Here, we show that CGK733 inhibits cell-cycle progression and global protein translation. Affinity purification identified adenine nucleotide translocator 2 (ANT2) as the primary target. CGK733 blocked ATP export from mitochondria and induced proton leak, thereby shifting ATP production from mitochondrial respiration to glycolysis and perturbing the TCA cycle. These mitochondrial alterations were accompanied by inactivation of the mTOR pathway and mild activation of the integrated stress response, resulting in translational inhibition. Collectively, our findings demonstrate that CGK733 acts mainly through ANT2-dependent mitochondrial modulation, revealing a mechanistic link between mitochondrial bioenergetics and translational control.

## Introduction

Chemical genetics has emerged as a powerful strategy for dissecting complex biological mechanisms using small molecules that acutely and reversibly modulate protein function. A critical step in this approach is to pinpoint the molecular target underlying a compound’s biological activity, as identifying the true target often uncovers previously unrecognized regulatory mechanisms and therapeutic opportunities. CGK733 was originally described as a potent inhibitor of ATM and ATR that suppresses cellular senescence^1^; however, that report was later retracted. Nevertheless, CGK733 continues to be marketed and cited as an ATM/ATR inhibitor^2–5^, whereas another study found no inhibitory activity toward either kinase^6^. Consequently, its molecular target and mechanism of action remain unresolved. Notably, CGK733 has been reported to inhibit cell proliferation, modulate tumor cell viability, induce endoplasmic reticulum (ER) stress, and regulate osteoclast differentiation^7–11^. Therefore, identifying the genuine molecular target of CGK733 would be of great value to researchers across multiple fields.

Many fundamental biological processes, including cell proliferation, stress responses, and differentiation, are closely linked to protein translation, which is a process indispensable for gene expression. Protein translation is one of the most ATP-consuming processes in eukaryotic cells^12,13^. Owing to its high energy demand, translation is tightly coupled to mitochondrial function, as mitochondria are the primary sites of ATP synthesis^14,15^. The mitochondrial respiratory chain pumps protons from the matrix into the intermembrane space, thereby generating the mitochondrial membrane potential^16,17^. ATP synthase, embedded in the inner mitochondrial membrane, utilizes this potential to drive ATP synthesis while allowing protons to re-enter the matrix^14,15^. However, not all of the pumped protons are coupled to ATP production; a fraction returns to the matrix without ATP synthesis, a process known as proton leak^17^. Given the high energy requirement of translation, energy-sensing pathways such as mTOR and AMPK regulate translational activity^18,19^. In particular, intracellular amino acid and ATP levels modulate mTOR activity, which in turn controls the phosphorylation of its downstream substrates and adjusts translational output.

Here, we sought to identify the molecular target and mechanism of action of CGK733, with a particular focus on how CGK733 inhibits cell proliferation and the underlying cellular processes.

## Results

### CGK733 affects both cell cycle progression and cell viability

We and others have shown that CGK733 (Fig. 1a) inhibits cell proliferation^7–9^; however, the detailed molecular mechanisms underlying this inhibition, as well as the molecular target of CGK733, remain unclear. To gain further insight into the molecular basis of this effect, we examined in detail the impact of CGK733 on cell proliferation. Treatment with ≥ 5 μM CGK733 markedly inhibited cell proliferation (Fig. 1b, left), whereas 10 μM or 20 μM CGK733 induced cell death (Fig. 1b, right). These findings suggest that 5 μM CGK733 inhibits cell-cycle progression without causing overt cytotoxicity. To test this hypothesis, we synchronized cells by a double-thymidine block and monitored the cell-cycle progression. As expected, 5 μM CGK733 delayed cell-cycle progression (Fig. 1c). Toghether, these results indicate that CGK733 affects both cell-cycle progression and cell viability in a concentration-dependent manner.

**Figure 1.**
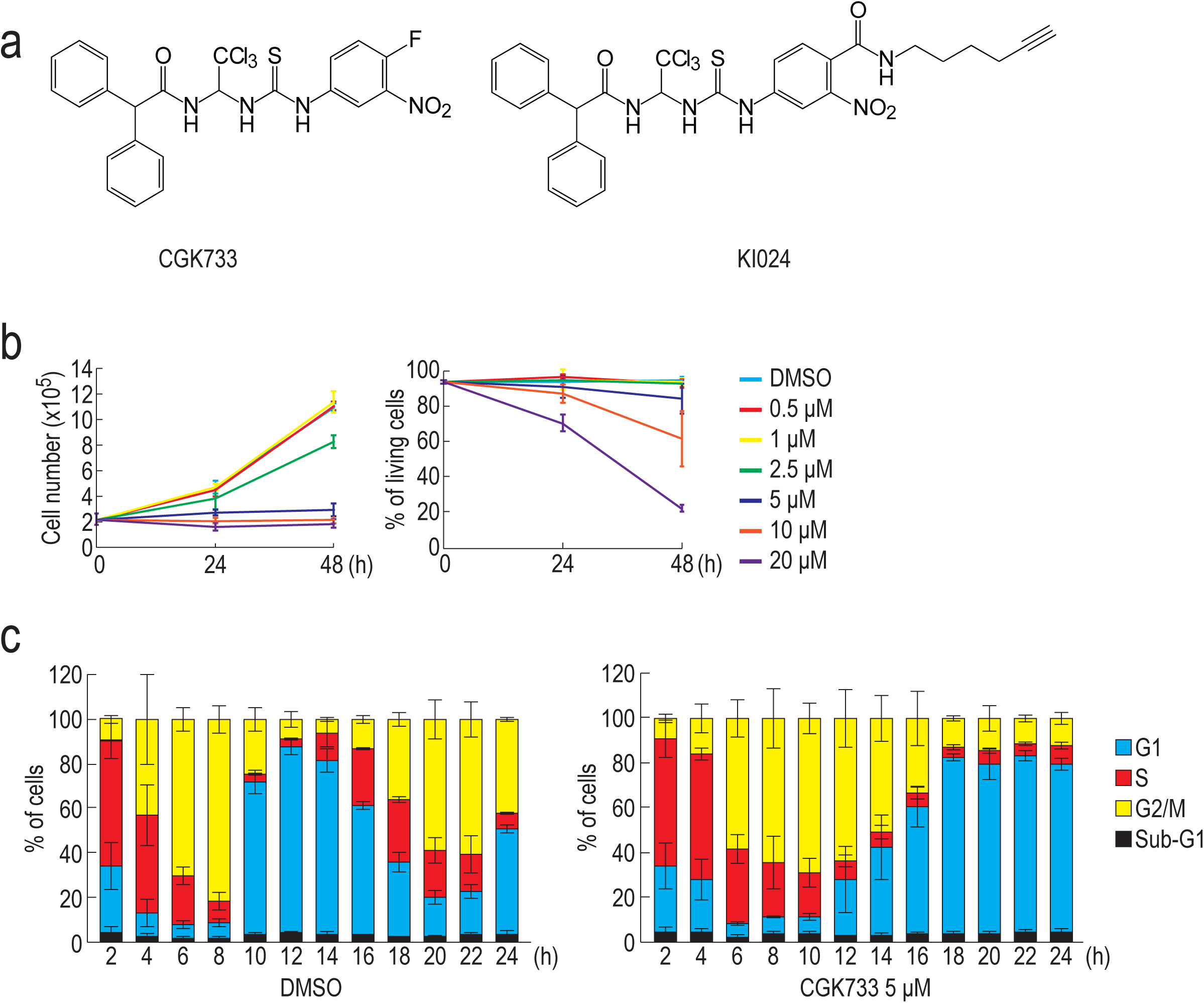
CGK733 suppresses cell cycle progression. (a) Chemical structures of CGK733 and KI-024. (b) HeLa cells were treated with the indicated concentrations of CGK733 for 24 or 48 h. Cell count and viability were assessed at each time point (n = 3). (c) Two hours after release from a double thymidine block, synchronized HeLa cells were treated with either DMSO or 5 μM CGK733. The cell cycle distribution was analyzed at the indicated time points using an image-based cytometer (n = 3). Error bars represent s.d.

### CGK733 inhibits global translation

To elucidate why CGK733 delays cell-cycle progression, we examined cyclin protein and mRNA levels after CGK733 treatment. A 6-h treatment with CGK733 reduced cyclin protein levels in a concentration-dependent manner with only a slight effect on PARP levels (Fig. 2a). In contrast, mRNA levels of cyclin A2, cyclin B1, and cyclin D3 remained unchanged after 6 h (Fig. 2b, upper). Under the same conditions, cyclin E1 mRNA levels decreased slightly, but its protein reduction was more pronounced. These results suggest that CGK733 affects either protein translation or degradation. A 24-h treatment with CGK733 also reduced cyclin protein levels in a concentration-dependent manner (Fig. 2a). Furthermore, treatment with 10 μM or 20 μM CGK733 for 24 h induced PARP cleavage – a hallmark of apoptosis^20^ – indicating that higher concentrations of CGK733 trigger apoptotic cell death, consistent with the results shown in Fig. 1b (Fig. 2a). Cyclin mRNA levels decreased slightly after treatment with low concentrations of CGK733 for 24 h, whereas 20 μM CGK733 markedly decreased them, presumably due to apoptotic cell death (Figs. 2a, b).

**Figure 2.**
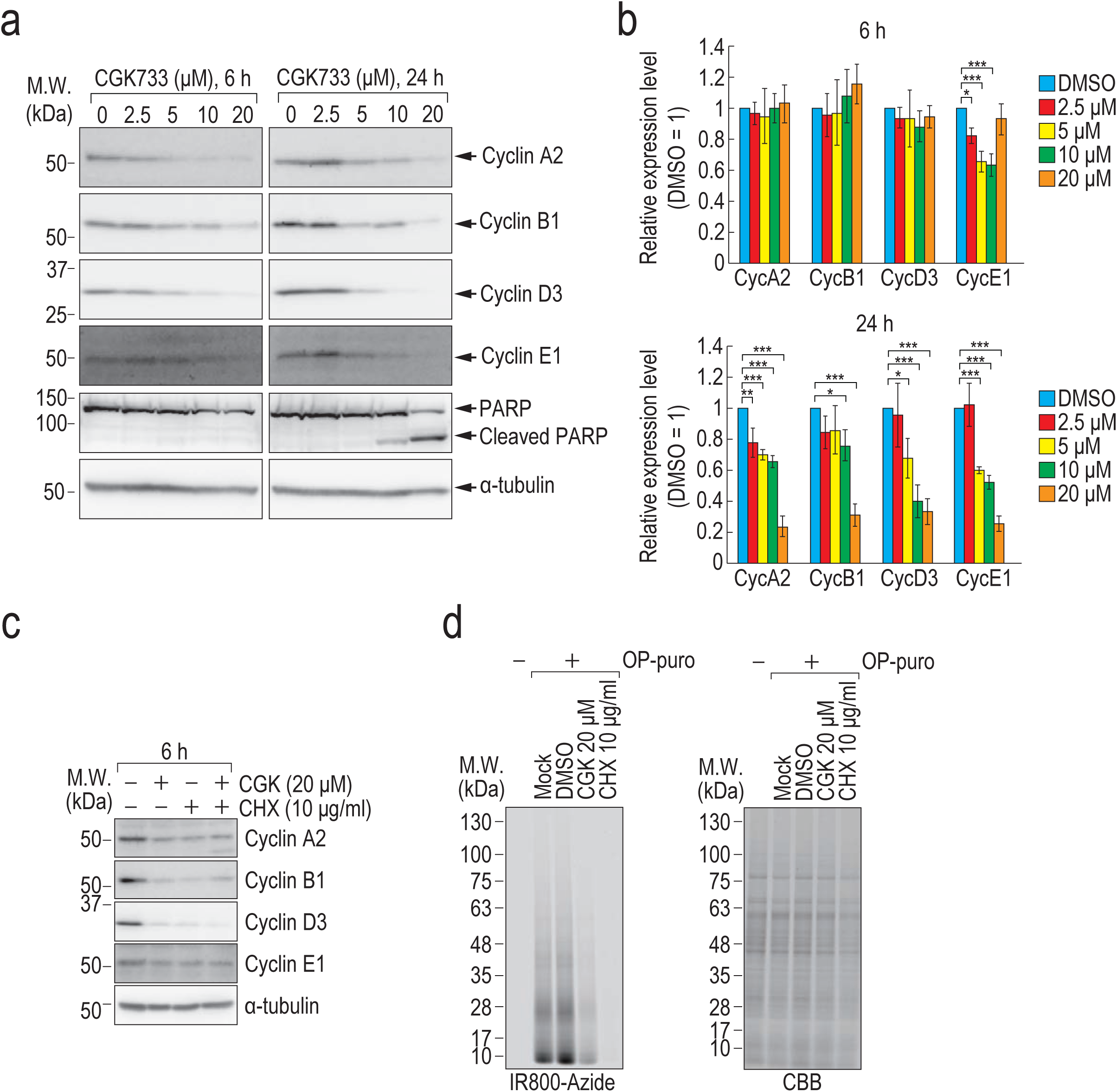
CGK733 inhibits protein translation. (a, b) HeLa cells were treated with the indicated concentrations of CGK733 for 6 or 24 h. Protein levels were analyzed by immunoblotting with the indicated antibodies (a), and RNA levels were measured by qRT-PCR (n = 3) (b). Statistical significance was determined by one-way ANOVA followed by Dunnett’s test (*: p < 0.05; **: p < 0.01; ***: p < 0.001). Error bars represent s.d. (c) HeLa cells were treated with 10 μM CGK733 and/or 10 μg/ml cycloheximide (CHX) for 6 h. Protein levels were analyzed by immunoblotting with the indicated antibodies. (d) HeLa cells were treated with 20 μM CGK733 for 3.5 h, followed by a 30 min-OP-puro treatment to label newly synthesized proteins. OP-puro-incorporated proteins were conjugated with Alexa Fluor 488 azide via a click reaction and detected using a fluorescence imaging system (left panel). Total protein was visualized by Coomassie Brilliant Blue staining (right panel).

Because CGK733 decreased cyclin protein levels without markedly affecting their mRNA levels, it is likely that CGK733 either promotes protein degradation or inhibits translation. Cyclin proteins are well known to undergo proteasomal degradation^21^, and a previous report suggested that CGK733 induces proteasome-dependent degradation of cyclin D1^7^. We therefore examined whether CGK733 enhances proteasomal degradation of cyclin proteins. However, CGK733 did not increase the abundance of proteasomal components (Extended Data Fig. 1a). We next investigated whether CGK733 affects the assembly of the 26S proteasome–the active form of the proteasome^22^–using native PAGE. Immunoblotting with an antibody against PSMA3 detected two upper bands and one lower band, whereas PSMC2 immunoblotting detected only the upper two bands (Extended Data Fig. 1b). Because PSMA3 is a 20S core-particle component and PSMC2 is a 19S regulatory-particle component, the upper two bands likely correspond to the 30S proteasome (one 20S core with two 19S particles) and the 26S proteasome (one 20S core with one 19S particle), while the lower band represents the free 20S proteasome. CGK733 treatment did not alter the distribution of these species (Extended Data Fig. 1b). Moreover, CGK733 had no effect on *in vitro* proteasome activity (Extended Data Fig. 1c). Alao and Sunnerhagen previously proposed that CGK733 promotes proteasomal degradation of cyclin D1, as the proteasome inhibitor MG132 restored cyclin D1 levels reduced by CGK733^7^. However, we also observed MG132-mediated restoration of cyclin levels in cells treated with the translation inhibitor cycloheximide (CHX) (Extended Data Fig. 1d). Thus, the MG132 effect likely reflects suppression of general protein turnover rather than reversal of a specific proteasome activation by CGK733. Collectively, these results indicate that CGK733 does not promote proteasomal degradation of cyclins.

We next tested the alternative hypothesis that CGK733 inhibits cyclin protein translation. Cyclin levels were examined in cells treated with CGK733 and/or CHX (Fig. 2c). Both CGK733 and CHX reduced cyclin protein levels to a similar extent, and their combined treatment produced no additive effect, suggesting that CGK733 inhibits cyclin protein synthesis. To directly assess whether CGK733 suppresses translation, we measured nascent protein synthesis using *O*-propargyl-puromycin (OP-puro) labelling^23^. In mock- or DMSO-treated cells, OP-puro-labeled nascent proteins were readily detected (Fig. 2d, left). In contrast, treatment with CGK733 or CHX markedly reduced OP-puro incorporation without affecting total protein levels (Fig. 2d). These results demonstrate that CGK733 suppresses global translation, similar to CHX.

### CGK733 binds to ANT2

As described above, CGK733 inhibits both cell proliferation and global translation (Figs. 1 and 2). However, CHX treatment, while inhibiting translation, did not induce cell death as CGK733 did (Extended Data Fig. 2), suggesting that translation inhibition alone cannot account for the cytotoxic effect of CGK733. Therefore, CGK733 likely inhibits translation indirectly with additional adverse outcomes. To clarify its mode of action, we sought to identify CGK733-binding proteins. Our previous structure-activity relationship study revealed that replacing the fluoro group of CGK733 with a bulkier substituent did not affect its antiproliferative activity^8^. Based on this finding, we synthesized a new probe, KI-024, to identify CGK733-binding proteins (Fig. 1a). To determine whether KI-024 retained the cell proliferation-inhibitory activity of CGK733, we compared their effects on cell growth. KI-024 inhibited cell proliferation, although its potency was approximately 50% that of CGK733 (Fig. 3a). Additionally, KI-024 treatment decreased cyclin A2 and B1 protein levels (Fig. 3b). These results indicate that KI-024 retains the ability to bind to the target protein responsible for the effects of CGK733 on cell proliferation and protein translation.

**Figure 3.**
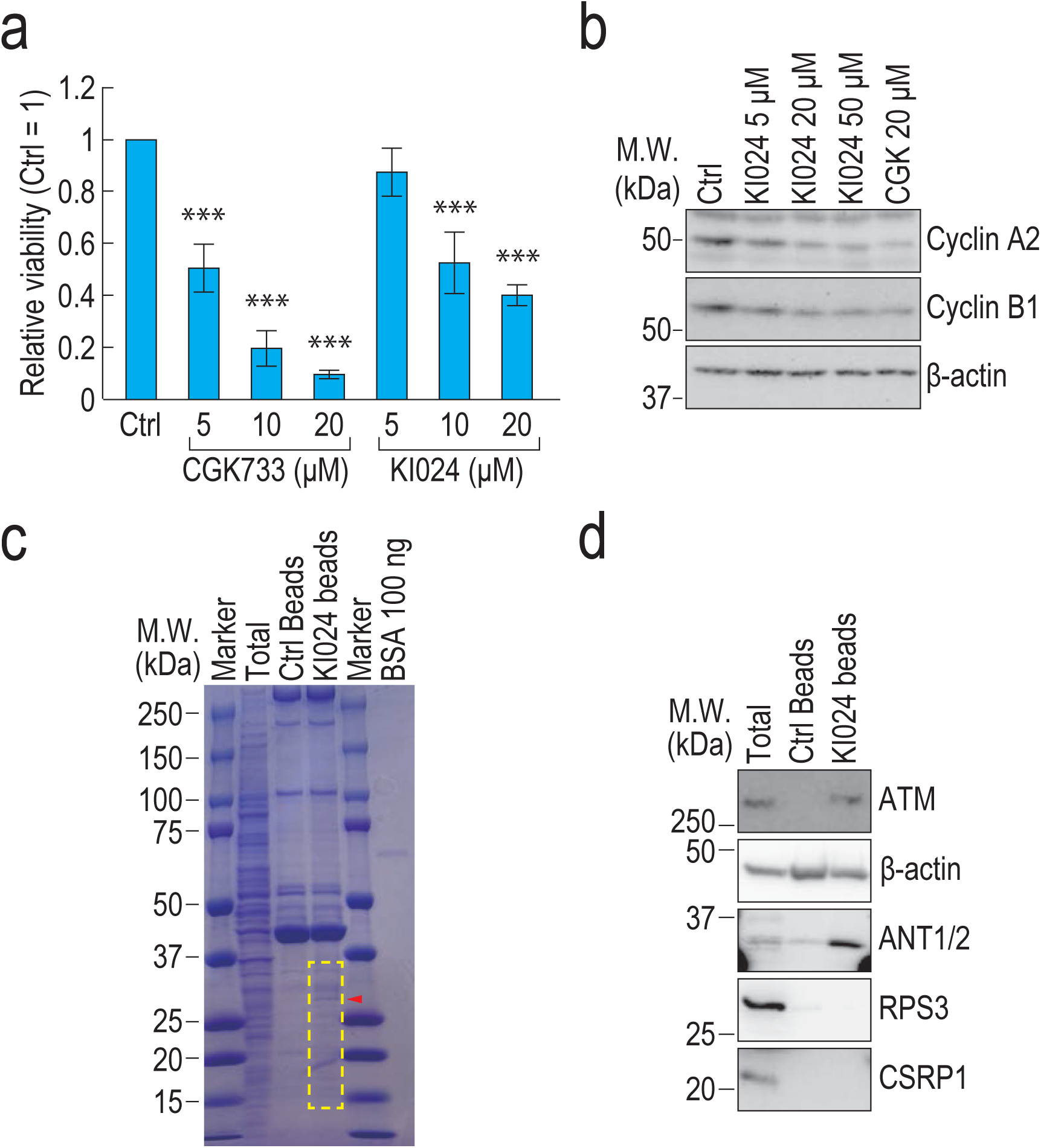
CGK733 binds to a mitochondrial ATP/ADP antiporter. (a) HeLa cells were treated with the indicated concentrations of CGK733 or KI-024 for 48 h, and cell viability was assessed (n = 3). Statistical significance was determined by one-way ANOVA followed by Dunnett’s test (***: p < 0.001). Error bars represent s.d. (b) HeLa cells were treated with the indicated concentrations of CGK733 or KI-024 for 6 h. Protein levels were analyzed by immunoblotting with the indicated antibodies. (c, d) CGK733-associated proteins were purified using KI-024-conjugated beads and subjected to SDS-PAGE followed by Coomassie Brilliant Blue staining. The red triangle indicates a KI-024-specific protein band of approximately 30 kDa, and the yellow rectangle marks the region that was excised and analyzed by LC-MS/MS (c) or analyzed by immunoblotting with the indicated antibodies (d).

Next, CGK733-binding proteins were purified using KI-024-conjugated magnetic beads and analyzed by SDS-PAGE followed by Coomassie Brilliant Blue staining (Fig. 3c). A prominent KI-024-specific band of approximately 30 kDa, along with several weaker bands, was detected and subjected to mass spectrometric analysis, which identified more than 60 candidate proteins (Extended Data Table 1). Based on sequence coverage and peptide counts, four proteins were selected for further validation. Among them, adenine nucleotide translocator 2 (ANT2) specifically interacted with KI-024-conjugated beads, as confirmed by immunoblotting (Fig. 3d), whereas β-actin, RPS3, and CSRP1 showed no specific binding. Collectively, these results indicate that ANT2 is the primary molecular target of CGK733.

Interestingly, we also detected an interaction between KI-024 conjugated beads and ATM, which had been previously reported as a CGK733-binding protein^1^. However, in this study, CGK733 did not inhibit the kinase activity of ATM (Extended Data Fig. 3). Etoposide, a topoisomerase II inhibitor that induces DNA double-strand breaks, and hydroxyurea, a ribonucleotide reductase inhibitor that impairs DNA replication, both activated the DNA damage response, leading to phosphorylation of ATM and Chk1/Chk2^24^ (Extended Data Fig. 3). The dual ATM/ATR inhibitor VE-821 and the ATM-specific inhibitor KU55933 suppressed the phosphorylation of ATM and Chk1/Chk2, whereas CGK733 had no effect on their phosphorylation status (Extended Data Fig. 3), consistent with a previous report^6^. These results indicate that, although CGK733 binds to ATM, it likely associates with a region not involved in its kinase activity.

### CGK733 causes accumulation of ATP in the mitochondrial matrix

ANT2 functions as a mitochondrial adenine nucleotide transporter that exports ATP synthesized in the mitochondrial matrix and imports ADP from the cytoplasm^25^. If CGK733 inhibits ANT2-mediated ATP export, ATP should accumulate within the matrix. To test this hypothesis, we employed a mitochondria-localized Förster resonance energy transfer (FRET)-based ATP reporter^26^. Treatment with 20 μM CGK733 increased reporter fluorescence at both 0.5 h and 4 h after treatment (Fig. 4a-c), suggesting ATP accumulation in mitochondria. Lower concentrations (1 μM and 5 μM) also caused transient ATP accumulation at 0.5 h, which returned to baseline by 4 h (Fig. 4a-c). These results indicate that CGK733 inhibits ANT2-mediated ATP export in a concentration-dependent manner. Notably, CGK733 treatment induced mitochondrial fragmentation–a hallmark of mitochondrial stress (Fig. 4a and Extended Data Fig. 4)–whereas it had no effect on the morphology of the ER, further supporting the notion that CGK733 primarily affects mitochondrial function.

**Figure 4.**
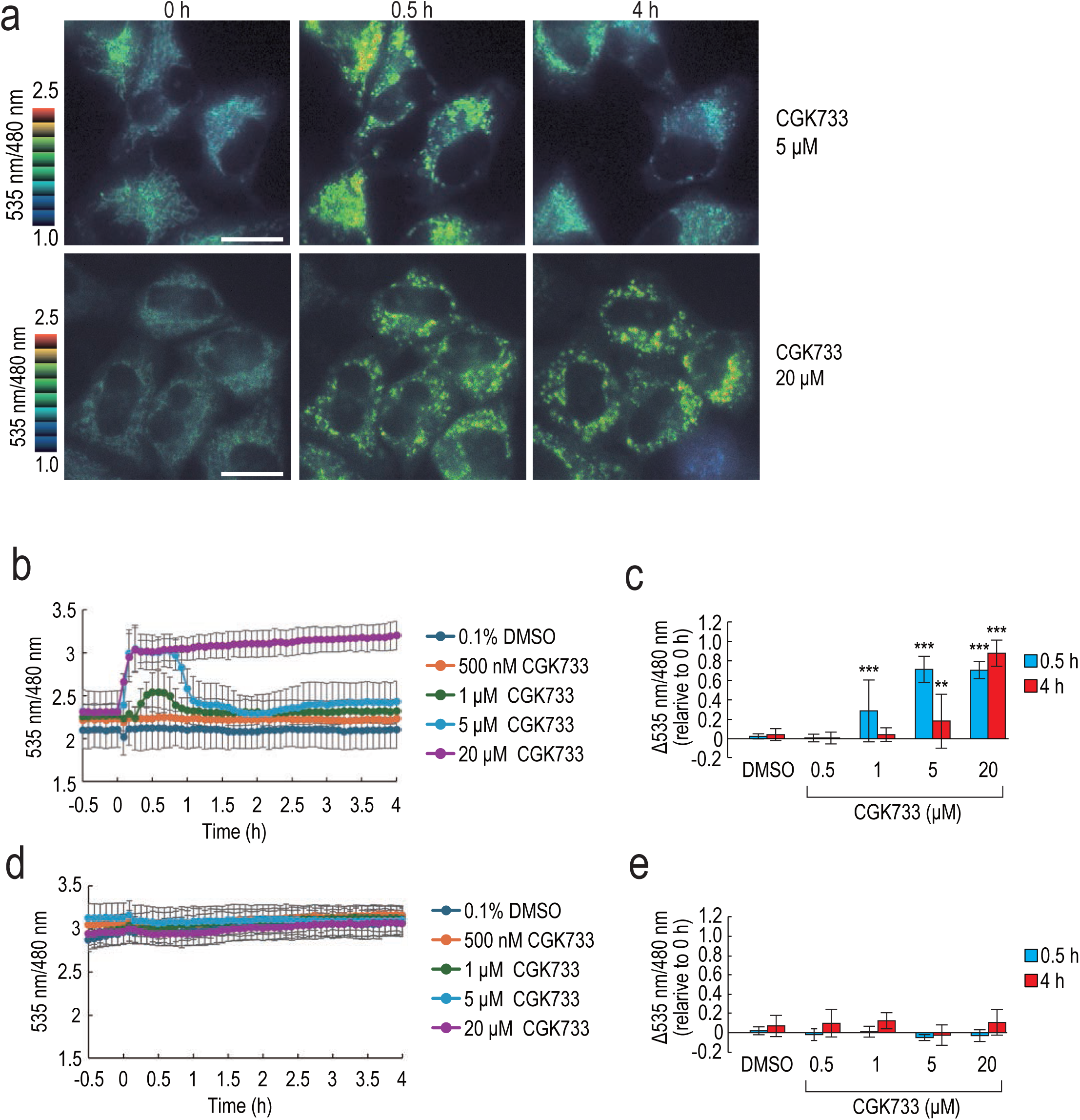
CGK733 increases mitochondrial ATP levels. (a) HeLa cells expressing mitATeam were treated with 5 or 20 μM CGK733 for 0.5 or 4 h, and fluorescence was observed. Pseudocolor images of mitochondrial emission ratio were shown. Scale bar, 10 µm. (b) HeLa cells expressing mitATeam were treated with 500 nM (n = 25), 1 µM (n = 27), 5 µM (n = 24), or 20 µM (n = 33) CGK733 or 0.1% DMSO (n = 19), and fluorescence was observed every 5 min. The averaged mitochondrial emission ratio was calculated. (c) HeLa cells expressing mitATeam were treated with 500 nM (n = 31 at 0.5 h; n = 30 at 4 h), 1 µM (n = 30 at 0.5 h; n = 29 at 4 h), 5 µM (n = 27 at 0.5 h; n = 27 at 4 h), or 20 µM (n = 36 at 0.5 h; n = 36 at 4 h) CGK733 or with 0.1% DMSO (n = 21 at 0.5 h; n = 22 at 4 h) for 0.5 or 4 h. Changes in mitochondrial emission ratios relative to 0 h were calculated. (d) HeLa cells expressing ATeam were treated with 500 nM (n = 27), 1 µM (n = 27), 5 µM (n = 30), or 20 µM (n = 33) CGK733 or 0.1% DMSO (n = 36), and fluorescence was observed every 5 min. The averaged cytoplasmic emission ratio was calculated. (e) HeLa cells expressing ATeam were treated with 500 nM (n = 27 at 0.5 h; n = 27 at 4 h), 1 µM (n = 29 at 0.5 h; n = 29 at 4 h), 5 µM (n = 28 at 0.5 h; n = 30 at 4 h), or 20 µM (n = 30 at 0.5 h; n = 33 at 4 h) CGK733 or with 0.1% DMSO (n = 35 at 0.5 h; n = 36 at 4 h) for 0.5 or 4 h. Changes in mitochondrial emission ratios relative to 0 h were calculated. Statistical significance was determined by one-way ANOVA followed by Dunnett’s test (**: p<0.01; ***: p<0.001). Error bars represent s.d.

We next monitored cytoplasmic ATP levels using a cytoplasm-localized FRET-based ATP reporter^26^, since inhibition of mitochondrial ATP export could potentially reduce cytosolic ATP. However, CGK733 did not significantly alter cytoplasmic ATP levels (Fig. 4d, e), presumably because glycolytic ATP production compensated for the reduced mitochondrial ATP export (see below for details).

### CGK733 stimulates proton leak

It has been reported that ANT2 also contributes to proton leak^27,28^, which refers to the movement of protons across the mitochondrial inner membrane without ATP synthesis^17^. To examine whether CGK733 affects this process, we measured the oxygen consumption rate (OCR)–an indicator of mitochondrial respiratory chain activity–after CGK733 treatment. CGK733 increased OCR, indicating enhanced proton leak and elevated oxygen consumption to maintain the mitochondrial membrane potential (Fig. 5a). Treatment with the ATP synthase inhibitor oligomycin decreased OCR in control cells, presumably because oxygen consumption coupled to ATP synthesis was abolished. In contrast, oligomycin had little effect on OCR in cells treated with 5 μM or 20 μM CGK733, suggesting that oxygen was consumed for proton leak rather than for ATP synthesis in these cells (Fig. 5a). Concomitantly, the extracellular acidification rate (ECAR)–an indicator of glycolytic activity–was increased in CGK733-treated cells, suggesting that glycolytic ATP production compensated for reduced mitochondrial ATP synthesis in these cells. Oligomycin did not further increase ECAR in CGK733-treated cells, consistent with the notion that oxygen consumption in these cells primarily reflected proton leak.

**Figure 5.**
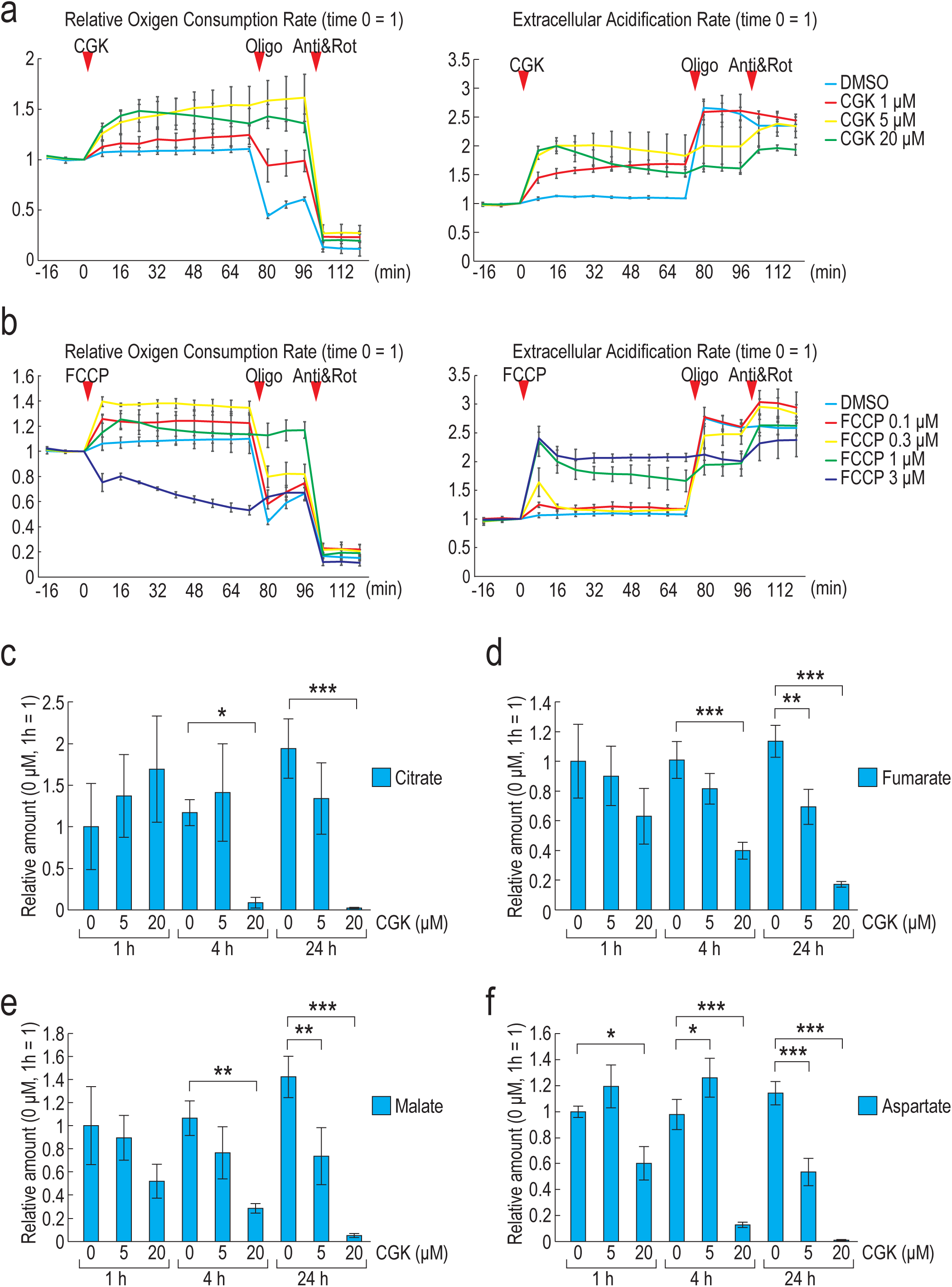
CGK733 induces proton leak. (a) Oxygen consumption rate (OCR) and extracellular acidification rate (ECAR) of HeLa cells were measured after sequential treatment with the indicated concentrations of CGK733, 1.5 μM oligomycin (oligo), and 0.5 μM rotenone/antimycin A (Rot/Anti) (n = 3). (b) OCR and ECAR were measured after sequential treatment with the indicated concentrations of FCCP, oligomycin and rotenone/antimycin A (n = 3). (c-f) The relative amounts of TCA cycle metabolites and aspartate in HeLa cells treated with the indicated concentrations of CGK733 for 1, 4, or 24 h (n = 3). Statistical significance was determined by one-way ANOVA and Dunnett’s test (*: p<0.05; **: p<0.01; ***: p<0.001). Error bars represent s.d.

From OCR and ECAR measurements, we calculated mitochondrial and glycolytic ATP production rates (Extended Data Fig. 5). CGK733 treatment decreased mitochondrial ATP production while increasing glycolytic ATP production, resulting in a modest reduction in total cellular ATP production (Extended Data Fig. 5). These results demonstrate that CGK733 shifts cellular energy metabolism from mitochondrial respiration to glycolysis, consistent with the observations described above (Figs. 4d, 4e and 5a). Treatment with FCCP, a well-known protonophore, mimicked the effects of CGK733 on OCR and ECAR, supporting the notion that CGK733 induces proton leak (Fig. 5b). Interestingly, 3 μM FCCP reduced OCR, presumably due to substrate exhaustion or excessive mitochondrial stress (Fig. 5b).

Because CGK733 induces proton leak and inhibits mitochondrial ATP production, we hypothesized that it might also affect the tricarboxylic acid (TCA) cycle, given the close link between mitochondrial respiration and TCA cycle flux^29^. Consistent with this, several TCA cycle metabolites, including malate, fumarate, and citrate, were significantly decreased after 4 h and 24 h of treatment with 20 μM CGK733, whereas some metabolites were increased, indicating altered TCA cycle flux (Fig. 5c-e and Extended Data Fig. 6). We also measured TCA cycle-related amino acids, including aspartate and glutamate, asparagine, and glutamine. Aspartate is derived from oxaloacetate and serves as a precursor for asparagine, whereas glutamate and glutamine are closely linked to α-ketoglutarate metabolism^29^. Notably, aspartate levels were markedly decreased following 4 h and 24 h of treatment with 20 μM CGK733, while asparagine levels were increased in cells treated with 5 μM CGK733 for 4 h and 24 h, and with 20 μM CGK733 for 4 h (Fig. 5f and Extended Data Fig. 6). Glutamine and glutamate levels were also decreased after 24 h of treatment with 20 μM CGK733. These results support the idea that CGK733 induces proton leak, thereby suppressing mitochondrial ATP production and perturbing TCA cycle activity.

Because proton leak generally reduces the mitochondrial membrane potential, we next monitored changes in membrane potential using the MT-1 dye, which emits red fluorescence when the potential is high. Unexpectedly, CGK733-treated cells retained red fluorescence, whereas FCCP-treated cells did not (Extended Data Fig. 7). We also used MitoTracker Red, which is another membrane potential-dependent dye. Consistent with the MT-1 result, CGK733-treated cells retained MitoTracker fluorescence (Extended Data Fig. 7). These results suggest that, although both CGK733 and FCCP induce proton leak, CGK733 does not collapse the mitochondrial membrane potential under our experimental conditions, by an as-yet unknown mechanism (See Discussion).

### CGK733 inactivates the mTOR pathway and activates the integrated stress response

As described above, CGK733 induces translational inhibition, suppresses mitochondrial ATP production, and decreases TCA cycle metabolites and aspartate levels (Figs. 2 and 5). Because the mTOR pathway is well known to sense intracellular ATP and amino acid levels^19^, we hypothesized that CGK733 inhibits translation through mTOR inactivation. To test this, we examined the phosphorylation status of p70S6K and 4E-BP1, both of which are well-established markers of mTOR activity^19^. As expected, CGK733 treatment caused dephosphorylation of p70S6K and 4E-BP1 at concentrations and time points similar to those that induced translation inhibition (Figs. 2 and 6a), suggesting that CGK733 inhibits translation via mTOR pathway inactivation.

**Figure 6.**
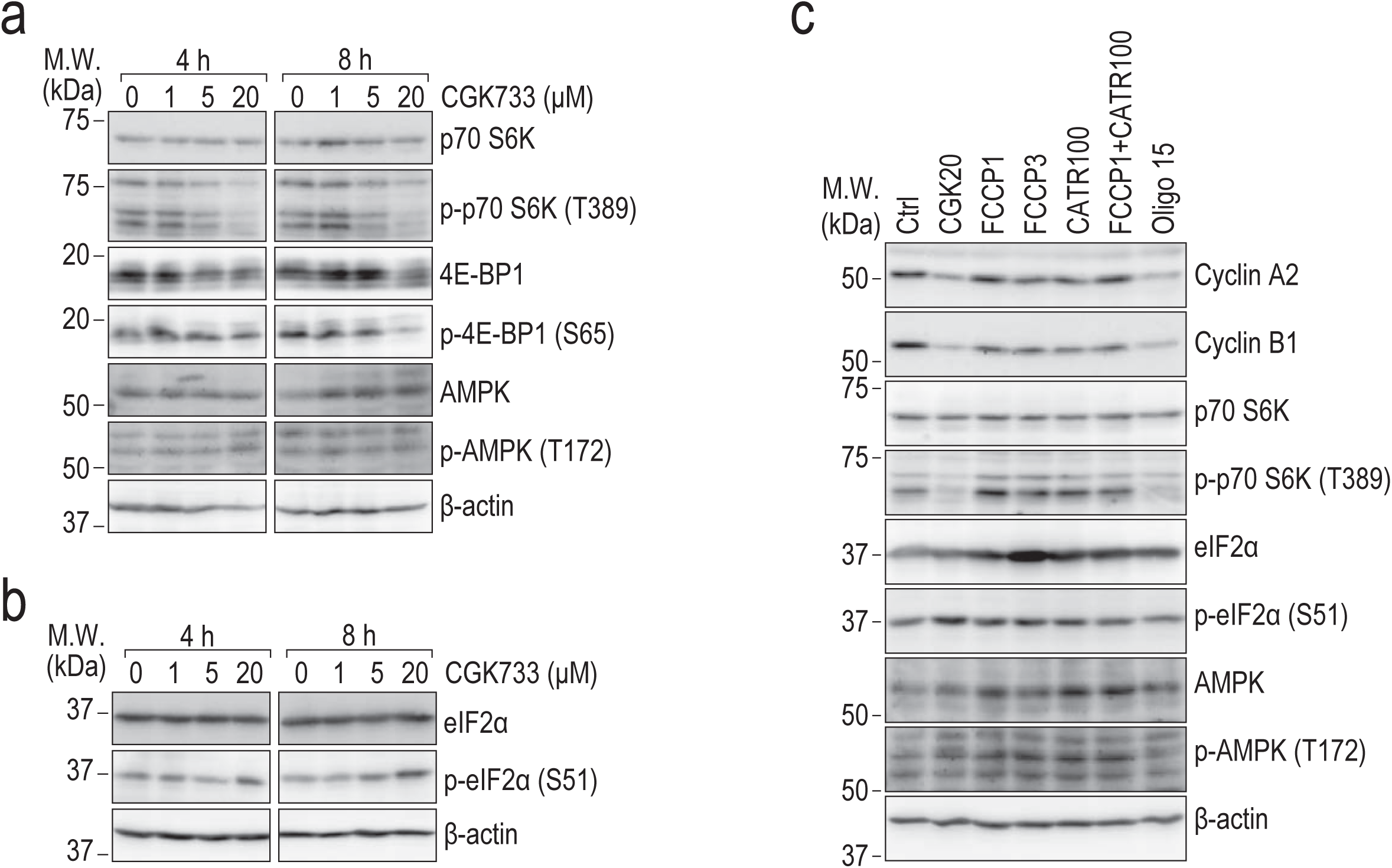
CGK733 inactivates the mTOR pathway. (a, b) HeLa cells were treated with the indicated concentrations of CGK733 for 4 or 8 h. Protein levels were analyzed by immunoblotting with the indicated antibodies. (c) HeLa cells were treated with 20 μM CGK733, 1 or 3 μM FCCP, 100 μM CATR, and 15 μM oligomycin for 6 h. Protein levels were analyzed by immunoblotting with the indicated antibodies.

AMP-activated protein kinase (AMPK) acts as a key sensor of cellular energy status^18^. When ATP levels decrease and the AMP/ATP (or ADP/ATP) ratio increases, AMPK becomes allosterically activated and phosphorylated. Activated AMPK then transduces signals to downstream pathways, including inhibition of the mTOR pathway. However, AMPK was not phosphorylated in CGK733-treated cells (Fig. 6a and Extended Data Fig. 5), suggesting that the modest reduction in total ATP production caused by CGK733 was insufficient to activate AMPK.

Mitochondrial stress and the accumulation of uncharged tRNAs are known to activate the integrated stress response (ISR) via distinct kinases such as HRI and GCN2, respectively, leading to phosphorylation of eIF2α and global translational inhibition^30^. Because CGK733 induces proton leak, inhibits ATP export, and causes mitochondrial morphological changes (Extended Data Fig. 4), CGK733 may trigger the ISR through mitochondrial stress. In addition, the decrease in aspartate levels may further activate the ISR by promoting the accumulation of uncharged tRNAs. To test these possibilities, we examined the phosphorylation status of eIF2α and found that CGK733 treatment slightly increased eIF2α phosphorylation (Fig. 6b), suggesting that ISR activation may partly contribute to the translational inhibition induced by CGK733. Collectively, these findings indicate that CGK733 inhibits translation through both mTOR pathway inactivation and mild ISR activation.

### Oligomycin treatment inhibits translation and inactivates the mTOR pathway

Finally, we sought to determine which of the biological activities of CGK733–namely, inhibition of ATP export, induction of proton leak, and suppression of mitochondrial ATP production–directly causes translational inhibition. To this end, we compared the effects of the ANT inhibitor carboxyatractyloside (CATR), the protonophore FCCP, and the ATP synthase inhibitor oligomycin on cyclin A2 and B1 translation. Although CGK733 inhibits ATP export and induces proton leak, neither individual treatment with CATR or FCCP nor their combined treatment affected cyclin A2 or B1 protein levels (Fig. 6c). However, because CATR did not induce ATP accumulation in mitochondria or alter the OCR (Extended Data Fig. 8), it is possible that CATR failed to reach mitochondria and inhibit ANT2 activity under our experimental conditions, presumably owing to its hydrophilicity. In contrast, oligomycin markedly reduced cyclin A2 and B1 protein levels (Fig. 6c), suggesting that inhibition of ATP synthase leads to translational inhibition. Oligomycin also caused dephosphorylation of p70S6K, indicating that oligomycin inactivates translation through the mTOR pathway (Fig. 6c). However, similar to CGK733, oligomycin did not induce phosphorylation of AMPK (Fig. 6c). Finally, we examined the phosphorylation status of eIF2α and found that oligomycin had no effect, in contrast to CGK733 (Fig. 6c).

Taken together, our findings demonstrate that CGK733 inhibits global translation, at least in part, through inactivation of the mTOR pathway and mild activation of the ISR. Because CGK733 binds to ANT2 and is associated with inhibition of ATP export, induction of proton leak, and suppression of mitochondrial ATP production, these results reveal a mechanistic link between mitochondrial function and translational control, providing new insights into how mitochondrial activity and cellular energy status regulate protein synthesis.

## Discussion

CGK733 is a compound with a controversial history: it was originally reported to inhibit ATM/ATR kinases and suppress cellular senescence, but the study was later retracted^1^. Consequently, its actual molecular target and mode of action have remained elusive. In this study, we found that CGK733 inhibits global translation, binds to ANT2, and modulates multiple mitochondrial functions, including ATP transport, proton leak, and mitochondrial ATP production.

We demonstrated that CGK733 suppresses global protein translation. However, unlike conventional translation inhibitors, CGK733 did not bind to translation factors such as ribosome and translation initiation factors^31^. In the CGK733-binding protein identification experiment, several ribosomal proteins were detected by mass spectrometry, but these proteins were most likely nonspecific contaminants owing to their high abundance. Indeed, no specific interaction between KI-024-conjugated beads and ribosomal protein RPS3 was detected by immunoblotting. These observations indicate that CGK733 inhibits translation indirectly rather than through direct interaction with the translation machinery.

We identified ANT2 as a CGK733 binding protein. To our knowledge, no prior study has reported a direct functional connection between ANT2 and the protein translation apparatus. However, because ANT2 regulates cellular energy metabolism, its dysfunction could indirectly affect translation through energy-sensing pathways such as mTOR and the ISR^19,25,30^. Indeed, CGK733 treatment led to marked inhibition of mTOR and mild activation of the ISR. The mTOR pathway is known to sense intracellular ATP and amino acid levels^19^. Although CGK733 slightly decreased total cellular ATP, the phosphorylation status of AMPK–an energy sensor that responds to cellular ATP concentration–remained unchanged, suggesting that the modest reduction in total ATP was insufficient to activate AMPK. Therefore, mTOR inactivation might instead result from reduced amino acid availability, particularly of aspartate. Several amino acid sensors for leucine, arginine, and methionine have been identified^19,32^, but no aspartate sensor has yet been reported. It is thus possible that an unidentified aspartate-sensing mechanism transmits this signal to mTOR.

Another intriguing possibility is that the mTOR pathway senses the activity or conformation of ATP synthase itself. Under our experimental conditions, CGK733, FCCP, and oligomycin did not alter AMPK phosphorylation, yet both CGK733 and oligomycin inactivated mTOR, whereas FCCP did not. These findings suggest that CGK733 and oligomycin inactivate mTOR through an AMPK-independent mechanism. Oligomycin binds to the F_0_ subunit of ATP synthase, blocking proton translocation and thereby inhibiting ATP synthesis^15^. CGK733 causes ATP accumulation and ADP depletion within the mitochondrial matrix, which may induce conformational and/or functional changes in ATP synthase similar to those caused by oligomycin. By contrast, FCCP dissipated the membrane potential, which may account for its different effects. Although the conformation of ATP synthase in CGK733-treated cells has not yet been determined, such analyses could provide valuable insight into how ATP synthase activity modulates mTOR signaling.

In addition to mTOR inactivation, CGK733 caused mild activation of the ISR. A previous study demonstrated that the ATP synthase inhibitor prethioviridamide activates the ISR by depleting ATP. Because ATP is essential for aminoacyl-tRNA synthesis, a decrease in ATP availability leads to the accumulation of uncharged tRNAs, thereby activating the ISR^33^. In contrast, in our experiments, CGK733 and FCCP induced a metabolic shift from mitochondrial respiration to glycolysis, resulting in only a slight reduction in total cellular ATP. Thus, these compounds (and presumably oligomycin) are unlikely to activate the ISR through ATP depletion under our experimental conditions. Indeed, neither FCCP nor oligomycin induced eIF2α phosphorylation, consistent with a previous report under different experimental conditions^34^. However, CGK733 did cause a modest increase in eIF2α phosphorylation. Because CGK733 has been reported to induce ER stress^10^, which can act upstream of ISR activation, it is plausible that CGK733 triggers mild ISR activation through ER stress under our experimental conditions.

As discussed above, CGK733 did not significantly alter mitochondrial membrane potential despite inducing proton leak. One plausible explanation is that its protonophoric activity was insufficient to cause detectable depolarization. Nevertheless, because the OCR pattern of CGK733-treatred cells closely resembled that of FCCP-treated cells, proton leak likely occurred. One possible explanation for the maintained membrane potential is that accumulation of negatively charged ATP⁴⁻ within the matrix compensates for the depolarization that would otherwise result from proton influx. Under normal conditions, the electrogenic ATP⁴⁻/ADP³⁻ exchange mediated by ANT exports one additional negative charge per transport cycle, slightly dissipating the membrane potential as an energetic cost of nucleotide transport^35,36^. When CGK733 inhibits this exchange, ATP⁴⁻ accumulates in the matrix, reducing dissipation through this pathway. The retained negative charge could counterbalance the proton influx caused by proton leak, thereby stabilizing the mitochondrial membrane potential in concert with the electron transport chain.

Another possible mechanism involves ATP hydrolysis-driven proton backflow. It has been reported that ATP synthase can hydrolyze ATP and pump protons from the matrix to the intermembrane space to maintain the membrane potential when it decreases^15^. This reverse activity has also been observed in FCCP-treated cells^37^. In CGK733-treated cells, matrix ATP accumulation may provide a more abundant substrate for this reverse reaction than in FCCP-treated cells, thereby facilitating reverse proton pumping and helping to maintain membrane potential. While these models are speculative, they highlight potential mechanisms by which CGK733 maintains mitochondrial polarization despite inducing proton leak. Further structural and biophysical studies will be required to clarify this mechanism.

In this study, we identified ANT2 as a direct binding partner of CGK733 and showed that CGK733 inhibits ATP export from the mitochondrial matrix. However, its binding mode remains unclear. CATR and another ANT inhibitor bongkrekic acid (BKA) are known to lock ANT in distinct conformations–the cytoplasmic-facing c-state and the matrix-facing m-state–thereby blocking nucleotide exchange^38–40^. CATR binds to the ATP/ADP translocation site from the cytoplasmic side and stabilizes the c-state, whereas BKA binds to an overlapping site within the same cavity and stabilizes the m-state. A recent structural study demonstrated that the binding sites of proton leak inducers such as FCCP and fatty acids overlap with those of ADP and CATR^27,38,41^. Considering that CGK733 both inhibits ATP transport and induces proton leak, it is plausible that CGK733 binds within the translocation cavity of ANT to mediate both effects simultaneously. Alternatively, given its hydrophobicity and membrane permeability, CGK733 might interact with the lipid-facing surface of ANT, stabilizing either the m- or c-state. Another possibility is that CGK733 functions as a hydrophobic protonophore, similar to FCCP, thereby promoting proton leak independently of direct ANT modulation^42^. Structural elucidation of the ANT2-CGK733 complex will be crucial to distinguish among these possibilities and to clarify the molecular basis of its dual activity.

A particularly interesting question regarding CGK733 is whether it inhibits ATM and ATR. In this study, we observed binding between CGK733 and ATM, consistent with the original report^1^, but CGK733 did not inhibit ATM or ATR signaling: phosphorylation of ATM and Chk1/Chk2 in response to DNA damage was unaffected. These findings, which are consistent with a previous report^6^, indicate that CGK733 is not an ATM/ATR inhibitor. Although the precise binding mode between ATM and CGK733 remains unknown, CGK733 may interact with a region unrelated to kinase activity.

Some earlier studies reported that CGK733 inhibits phosphorylation of RPA2 in XP30RO cells and p53 in IEC-6 cells after treatment with DNA-damaging agents^2,3^. Because these proteins are well-known ATM/ATR substrates^43–47^, CGK733 was interpreted as an ATM/ATR inhibitor. What might explain this discrepancy? IEC-6 and XP30RO are derived from normal rat intestinal epithelium and from primary dermal fibroblasts of a xeroderma pigmentosum variant patient, respectively, whereas our study used HeLa cells (cervical cancer), and Choi et al. used H460 cells (lung cancer).

Normal cells rely more heavily on mitochondrial respiration than glycolysis for ATP production^48,49^. Thus, their cellular ATP levels may be more sensitive to CGK733, which disrupts mitochondrial ATP synthesis. Because ATP depletion has been reported to reduce global protein phosphorylation^50,51^, differences in energy metabolism could underlie the cell type-specific (cancer vs non-cancer cells) effects of CGK733. However, metabolic profiles of IEC-6 and XP30RO cells have not been reported, while Wu et al. showed that H460 cells are more glycolytic than A549 lung cancer cells^52^.

Thus, the discrepancy likely reflects metabolic differences between cancer and non-cancer cells. While our data strongly argue against CGK733 being an ATM/ATR inhibitor, we cannot completely exclude the possibility that it affects ATM/ATR activity in a context-dependent manner. Further comprehensive analyses will be required to elucidate the molecular basis underlying these effects.

## Acknowledgments

We thank Mr. S. Ueno for technical assistance, Dr. H. Imamura (Yamaguchi University) for providing the ATeam and mitAteam plasmids, and Dr. K. Matsumoto for critical discussions. This work was supported in part by JSPS KAKENHI (Grant Numbers JP20K21247 and JP23K18418 to D.K.; JP23H05473 and JP23H04882 to M.Y.; JP21H02423 to S.I.); the Koyanagi Foundation; the Tamura Science & Technology Foundation; the Takahashi Industrial and Economic Research Foundation; the SENSHIN Medical Research Foundation; University of Toyama (President’s Discretionary Research Fund) (all to D.K.); and RIKEN (Pioneering Project and RIKEN TRIP initiative “TRIP-AGIS” to S.I.).

## Contributions

D.K. conceived the idea, conceptualized the project, and performed the cell proliferation assay, qRT-PCR, immunoblotting, isolation of KI-024-binding proteins, and Seahorse metabolic assay. Y.I, K.I., and R.K. synthesized KI-024. S.I. and M.S. performed the OP-puro assay. K.S. and M.Y. performed FRET analysis. K.Y. and T.N. performed metabolite measurements. D.K. and S.K.I. performed native PAGE. T.I. performed the proteasome activity assay. S.K. performed mitochondrial morphology observation and analysis of the mitochondrial membrane potential. D.K. wrote the manuscript, and all authors revised and approved the final version.

## Materials and Methods

### Cell culture and synchronization

HeLa S3 (RIKEN BRC Cell Bank, #RCB0191) and HEK293 Flp-In T-REx (Thermo Fisher Scientific, R78007) cells were cultured in Dulbecco’s modified Eagle’s medium (DMEM) supplemented with 10% heat-inactivated fetal bovine serum (Thermo Fisher Scientific, Waltham, MA, USA). Cells were maintained in 5% CO_2_ at 37°C. For cell-cycle synchronization, cells were treated with 2 mM thymidine (FUJIFILM Wako Pure Chemical Corporation, Osaka, Japan) for 18 h. After treatment, the cells were washed twice with phosphate-buffered saline (PBS) to release them from the thymidine block and cultured in fresh culture medium for 8 h. The cells were then treated again with 2 mM thymidine for 16 h and washed twice with PBS to release them from the double-thymidine block.

### Antibodies and reagents

Mouse monoclonal anti-α-tubulin antibody (T6074) was obtained from Sigma-Aldrich (St. Louis, MO, USA). The following antibodies were purchased from Cell Signaling Technology (Danvers, MA, USA): mouse monoclonal anti-cyclin A2 (#4656), rabbit polyclonal anti-cyclin B1 (#4138), mouse monoclonal anti-cyclin D3 (#2936), mouse monoclonal anti-cyclin E1 (#4129), rabbit monoclonal anti-ATM (#2873), rabbit monoclonal anti-phospho-ATM (S1981), mouse monoclonal anti-Chk1 (#2360), rabbit monoclonal anti-phospho-Chk1 (S345) (#2348), mouse monoclonal anti-Chk2 (#3440), and rabbit polyclonal anti-phospho-Chk2 (T68). Rabbit polyclonal anti-ANT1/2 (17796-1-AP), mouse monoclonal anti-RPS3 (66046-1-Ig), rabbit polyclonal anti-CSRP1 (13432-1-AP), and rabbit polyclonal anti-phospho-p70S6K (T389) (28735-1-AP) antibodies were obtained from Proteintech (Rosemont, IL, USA). Rabbit polyclonal anti-PARP-1 (sc-7150), mouse monoclonal anti-PSMA3 (sc-166205), mouse monoclonal anti-PSMB1 (sc-374405), and mouse monoclonal anti-PSMC2 (sc-166972) antibodies were obtained from Santa Cruz Biotechnology (Santa Cruz, CA, USA). Mouse monoclonal anti-β-actin antibody was purchased from Medical and Biological Laboratories (MBL, Nagoya, Japan). Rabbit monoclonal anti-p70S6K (F0011), rabbit monoclonal anti-4E-BP1 (F0128), rabbit monoclonal anti-phospho-4E-BP1 (S65) (F0339), rabbit monoclonal anti-AMPKα (F0269), rabbit monoclonal anti-phospho-AMPKα (T172) (F0151), rabbit monoclonal anti-eIF2α (F0316), and rabbit monoclonal anti-phospho-eIF2α (S51) (F0257) antibodies were purchased from Selleck Chemicals LLC (Houston, TX, USA). HRP-conjugated anti-mouse IgG and anti-rabbit IgG secondary antibodies were purchased from GE Healthcare (Chicago, IL, USA). CGK733 used in this study was synthesized as previously described^8^. Puromycin and G418 were purchased from Takara Bio (Otsu, Japan). MG132, etoposide, cycloheximide, and hydroxyurea were purchased from Sigma-Aldrich. Oligomycin was obtained from Cayman Chemical (Ann Arbor, MI, USA). FCCP was purchased from MedChemExpress (Monmouth Junction, NJ, USA). KU55933 and VE-821 were obtained from Adooq Bioscience (Irvine, CA, USA).

### Synthesis of KI-024

KI-024 was synthesized according to the procedure used for the synthesis of CGK733 and its derivatives, with different substrates^1,8^.

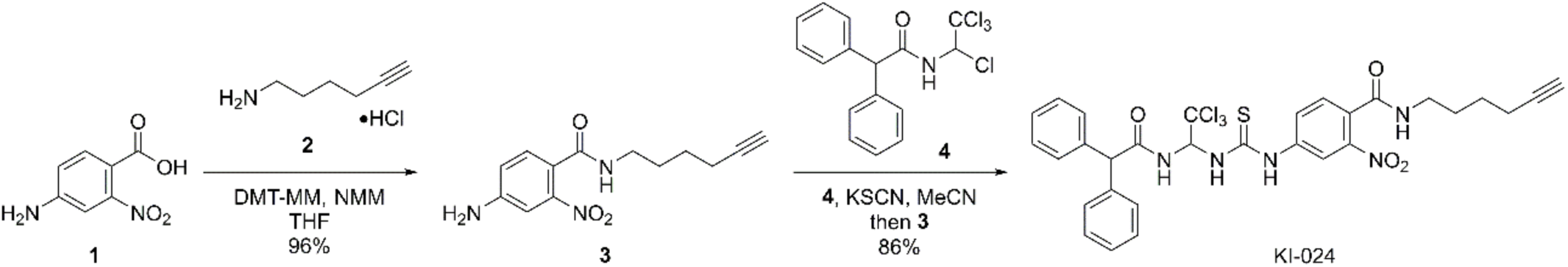

Infrared (IR) spectra were measured with a JASCO FT/IR-4100 spectrophotometer (JASCO, Tokyo, Japan) equipped with an ATR PRO ONE attachment. Nuclear magnetic resonance (NMR) spectra were recorded on a JEOL JNM-ECZ 400S/L1 spectrometer (JEOL, Tokyo, Japan). Chemical shifts (δ, ppm) were referenced to the residual solvent peaks as the internal standards (DMSO-*d*_6_: δ_H_ = 2.49, δ_c_ = 39.5). Mass spectra were recorded on a JEOL JMS-T100LP spectrometer. All reactions were carried out under an argon atmosphere.

To a stirred solution of 4-amino-2-nitrobenzoic acid (**1**, 273 mg, 1.50 mmol) and hex-5-yn-1-amine hydrochloride (**2**, 241 mg, 1.80 mmol) in tetrahydrofuran (THF, 15 mL) were added 4-(4,6-dimethoxy-1,3,5-triazin-2-yl)-4-methylmorpholinium chloride (DMT-MM, 498 mg, 1.80 mmol) and *N*-methylmorpholine (NMM, 412 µL, 3.75 mmol). The reaction mixture was stirred at room temperature for 5 h, then concentrated under reduced pressure. The residue was dissolved in EtOAc, washed successively with water and brine, dried over anhydrous Na_2_SO_4_, and concentrated under reduced pressure. The residue was passed through silica gel (eluent: CHCl_3_/MeOH = 40:1) and subsequently recrystallized from MeOH/EtOAc/hexane to yield 4-amino-*N*-(hex-5-yn-1-yl)-2-nitrobenzamide (**3**, 376 mg, 96%) as a yellow solid.

Compound **5**: IR (ATR): ν_max_ (cm^-1^) = 3429, 3287, 3073, 2950, 2904, 1629, 1524, 1457, 1433, 1364, 1310. ^1^H NMR (400 MHz, DMSO-*d*_6_): δ (ppm) = 1.42-1.58 (4H, m), 2.17 (2H, dt, *J* = 2.4, 6.8 Hz), 2.76 (1H, t, *J* = 2.4 Hz), 3.14 (2H, q, *J* = 6.4 Hz), 6.05 (2H, s), 6.74 (1H, dd, *J* = 8.4, 2.4 Hz), 6.95 (1H, d, *J* = 2.4 Hz), 7.25 (1H, d, *J* = 8.4 Hz), 8.32 (1H, t, *J* = 5.4 Hz). ^13^C NMR (100 MHz, DMSO-*d*_6_: δ (ppm) = 17.4, 25.4, 28.1, 38.4, 71.3, 84.4, 107.4, 115.8, 118.1, 129.9, 149.6, 150.9, 165.3. HRMS (ESI/TOF): *m/z* calcd. for C_13_H_15_N_3_O_3_Na [M+Na]^+^ 284.1006; found 284.1002.

KSCN (43.7 mg, 450 µmol) was added to a solution of 2,2-diphenyl-*N*-(1,2,2,2-tetrachloroethyl)acetamide^8^ (**4**, 113 mg, 300 µmol) in MeCN (3.0 mL), and the mixture was stirred at room temperature for 1 h. Compound **3** (78.4 mg, 300 μmol) was then added, and the reaction mixture was stirred at room temperature for 24 h. Water was added to the reaction mixture, and the resulting precipitate was collected by filtration. Reprecipitation from EtOAc/MeOH/MeCN afforded KI-024 (171 mg, 86%) as a slightly yellow solid.

KI-024: IR (ATR): ν_max_ (cm^-1^) = 3335, 3284, 3266, 3087, 2973, 2936, 2875, 1693, 1639, 1583, 1536, 1492, 1463, 1452, 1334, 1305, 1194, 1138, 1102, 1028. ^1^H NMR (400 MHz, DMSO-*d*_6_): δ (ppm) = 1.44-1.63 (4H, m), 2.19 (2H, dt, *J* = 2.4, 6.8 Hz), 2.77 (1H, t, *J* = 2.4 Hz), 3.20 (2H, q, *J* = 6.4 Hz), 5.15 (1H, s), 7.18-7.36 (11H, m), 7.54 (1H, d, *J* = 8.4 Hz), 7.84 (1H, dd, *J* = 8.4, 2.4 Hz), 8.42 (1H, d, *J* = 2.4 Hz), 8.44 (1H, d, *J* = 9.2 Hz), 8.64 (1H, t, *J* = 6.4 Hz), 9.23 (1H, d, *J* = 8.4 Hz), 10.70 (1H, s). ^13^C NMR (100 MHz, DMSO-*d*_6_: δ (ppm) = 17.4, 25.3, 27.9, 38.5, 56.1, 69.5, 71.3, 84.4, 101.1, 117.2, 126.3, 126.8, 126.8, 128.1, 128.3, 128.3, 128.6, 128.7, 129.3, 139.6, 139.8, 140.7, 146.9, 164.9, 170.6, 180.8. HRMS (ESI/TOF): *m/z* calcd. for C_30_H_28_N_5_O_4_NaSCl_3_ [M+Na]^+^ 682.0820; found 682.0820.

### Stable cell line establishment

To establish HeLa cells stably expressing ATeam or mitATeam, HeLa cells were transfected with pcDNA3.1-AT1.03 or pcDNA3.1-mitAT1.03^26^, and stable clones were selected by treatement with puromycin (1 µg/ml) followed by clonal isolation. Stable clones were maintained in medium containing puromycin and G418.

### Cell count and cell viability assay

After trypsinization, viable and dead cells were counted using the trypan blue exclusion method with a hemocytometer (Fig. 1). Cell viability was also assayed using the Cell Count Reagent SF (nacalai tesque, Kyoto, Japan) according to the manufacturer’s instructions (Fig. 3).

### Cell cycle analysis

Cells were fixed in 70% ethanol, rinsed with PBS, and then stained with a solution containing 20 μg/ml propidium iodide (Thermo Fisher Scientific), 0.05% Triton X-100, and 0.1 mg/ml RNase A (Thermo Fisher Scientific). The cell-cycle distribution was analyzed using a Tali image-based cytometer (Thermo Fisher Scientific).

### Immunoblotting

Cells were lysed directly on plates in 1× SDS-PAGE sample buffer. Proteins were then separated by SDS-PAGE and transferred onto a PVDF membranes by electroblotting. Membranes were incubated with primary and secondary antibodies using standard procedures, and protein bands were visualized using a NOVEX ECL Chemiluminescent Substrate Reagent Kit (Thermo Fisher Scientific) and an ImageQuant LAS 4000 mini system (GE Healthcare).

### RNA preparation and RT-PCR

Total RNA was extracted using TRIzol reagent (Thermo Fisher Scientific) according to the manufacturer’s instructions. cDNA was synthesized from total RNA using PrimeScript Reverse Transcriptase (Takara Bio) with random primers. Quantitative RT-PCR was performed with an MX3000P system (Agilent Technologies, Santa Clara, CA, USA) using SYBR Green chemistry, and relative mRNA levels were calculated using the comparative Ct method. mRNA levels were normalized to 18S rRNA. All primer sequences are listed in Table S1.

### Nascent peptide labeling with OP-puro

Nascent peptide labeling with OP-puro was performed as previously described with minor modifications^23^. Cells seeded in 24-well plates were treated with 20 μM CGK733 or 10 μg/ml CHX for 3.5 h, incubated with 20 μM OP-puro (Jena Bioscience) for 30 min, washed with 1 ml of phosphate-buffered saline (PBS), and lysed in 60 µl of OP-puro lysis buffer (20 mM Tris-HCl pH 7.5, 150 mM NaCl, 5 mM MgCl_2_, and 1% Triton X-100). After the removal of cell debris by centrifugation, lysates were incubated with 1 μM IRDye 800CW azide (LI-COR Biosciences) for 30 min at 25 °C using a Click-iT Cell Reaction Buffer Kit (Thermo Fisher Scientific). After the removal of unreacted azide by MicroSpin G-25 columns (Cytiva), the proteins were separated by sodium dodecyl sulfate (SDS)-polyacrylamide gel electrophoresis (PAGE). The infrared (IR) 800 signals were detected using an ODYSSEY CLx imaging system (LI-COR Biosciences). The total protein in the gel was stained with EzStain AQua (ATTO) and detected by ODYSSEY CLx (LI-COR Biosciences) with an IR700 channel. Total protein-normalized nascent peptide signals were quantified after background subtraction.

### Identification of CGK733 binding proteins

KI-024 was conjugated to azide beads (FG beads, Tamagawa Seiki, Nagano, Japan) according to the manufacturer’s instructions. HeLa cells were suspended in lysis buffer [25 mM HEPES (pH 7.5), 150 mM NaCl, 2 mM MgCl_2_, 1 mM EDTA, 2.5 mM EGTA, 1% Nonidet P-40, 10% glycerol, cOmplete Protease Inhibitor Cocktail (Sigma-Aldrich), and PhosSTOP (Sigma-Aldrich)] and sonicated for 10 s. After centrifugation, cell extracts were incubated with KI-024 conjugated beads for 4 h at 4 °C with gentle agitation. The beads were washed with lysis buffer three times and suspended in 1× SDS sample buffer,and incubated at 95 °C for 5 min. Protein samples were subjected to SDS-PAGE and stained with Coomassie Brilliant Blue. Bands corresponding to CGK733 binding proteins were excised and subjected to LC-MS/MS analysis (Japan Proteomics Co., Sendai, Japan).

### Förster resonance energy transfer (FRET)

HeLa cells stably expressing ATeam 1.03 or mitATeam 1.03 were cultured in DMEM without phenol red to improve imaging quality. Wide-field fluorescence imaging was performed on an IX83 microscope (Evident, Tokyo, Japan) equipped with an ORCA-FLASH4.0 CMOS camera (Hamamatsu Photonics, Hamamatsu, Japan) and a 60× objective lens (PlanApo N60× NA1.42 Oil) at 37 °C under 95% air and 5% CO_2_. Cells were illuminated with a 447 nm LED light source (X-Cite XLED1, Excelitas, Pittsburgh, PA, USA) through a 25% ND filter. Images were collected using MetaMorph software (Universal Imaging, Bedford Hills, NY, USA) with a 440AF21 excitation filter, a 455DRLP dichroic mirror, and two emission filters (480AF30 for CFP and 535AF26 for Venus).

### Measurement of OCR and ECAR

Oxygen consumption rate (OCR) and extracellular acidification rate (ECAR) were measured using an Agilent Seahorse XFe24 Extracellular Flux Analyzer with an Agilent Seahorse XF Real-Time ATP Rate Assay Kit (Agilent Technologies). Cells were seeded in XF24 cell culture microplates at a density of 3 × 10^4^ cells/well one day before the assay. On the day of measurement, the culture medium was replaced with Seahorse XF DMEM Medium (pH 7.4) supplemented with 10 mM glucose, 1 mM pyruvate, and 2 mM glutamine, and cells were incubated for 1 h at 37 °C in a non-CO₂ incubator. Basal OCR and ECAR were measured three times, followed by nine measurement cycles after addition of CGK733, FCCP, or CATR. Subsequently, OCR and ECAR were measured three times after oligomycin (1.5 μM) addition and three times after rotenone/antimycin A (0.5 μM each) addition. Each measurement cycle consisted of 3 min mixing, 2 min waiting, and 3 min measuring. Data were analyzed using Wave software (Agilent Technologies).

### Measurement of metabolites

Metabolite extraction and metabolomic analyses were performed as previously described^53–56^. Briefly, cells were treated with CGK733 for 1, 4, or 24 h and then collected in ice-cold 50% methanol/50% water. The homogenate was mixed with an equal volume of chloroform and vortexed. The mixture was centrifuged at 13,000 ×g for 10 min at 4 °C. The upper aqueous phase was collected, mixed again with chloroform, and centrifuged under the same conditions. The aqueous phase was then dried using a SpeedVac SPD1010 (Thermo Fisher Scientific).

For amino acid analysis, dried samples were reconstituted in LC-MS-grade water and derivatized with *N*^α^-(5-fluoro-2,4-dinitrophenyl)-L-leucinamide (L-FDLA). Each sample (50 µL) was mixed with 10 µL of 200 mM sodium bicarbonate and 10 µL of 1% L-FDLA in acetone, followed by incubation at 40 °C for 1 h. After dilution with 50% methanol, the resulting solution was filtered through a 0.45-µm membrane and subjected to LC–MS/MS analysis. Chromatographic separation was performed on an MG3 reversed-phase ODS column (2.0 × 150 mm, 3 µm; Osaka Soda) using mobile phase A (5 mM ammonium formate) and mobile phase B (methanol) at a flow rate of 150 µL/min and a column temperature of 40 °C. The programmed gradient was as follows: 0–10 min, 20–80% B; 10–15 min, 80% B; 15–20 min, 20% B. Metabolites were analyzed using an Agilent 6460 Triple Quadrupole mass spectrometer coupled to an Agilent 1290 HPLC system.

For GC-MS analysis, dried samples were derivatized with methoxyamine hydrochloride in pyridine at 30 °C for 90 min, followed by derivatization with N-methyl-N-trimethylsilyltrifluoroacetamide containing 1% trimethylchlorosilane (MSTFA + 1% TMCS; Pierce) at 37 °C for 30 min. TCA cycle intermediates were analyzed in selected ion monitoring (SIM) mode using an Agilent 5977 MSD single quadrupole mass spectrometer coupled to an Agilent 7890 gas chromatograph. Metabolite separation was achieved on a 30 m DB-5MS column with a 10 m DuraGuard precolumn. Helium was used as the carrier gas at a constant flow rate of 1.1 mL/min. The oven temperature program was as follows: initial hold at 60 °C for 1 min, ramp at 10 °C/min to 325 °C, and hold at 325 °C for 10 min. Metabolites were quantified by peak integration using the MassHunter Quantitative Analysis software (Agilent Technologies) and normalized to cell number.

### Proteasome activity assays

In vitro proteasome degradation assays were performed as previously described^57^. Briefly, 2 nM 26S yeast proteasome and 125 nM Suc-LLVY-AMC (Peptide Institute) in digestion buffer (50 mM Tris-HCl (pH 7.5), 5% (v/v) glycerol, 5 mM MgCl_2_, 1 mM DTT, 1 mM ATP, and 0.5% DMSO) were incubated at 30 °C. Emission fluorescence at 460 nm (excitation at 360 nm) was measured every 0.5 min for 30 min using a Synergy H1 plate reader (Agilent Technologies).

### Native gel electrophoresis

Native gel electrophoresis was performed essentially as described previously^58^, with minor modifications. Cells were washed with ice-cold PBS and lysed directly in lysis buffer containing 20 mM Tris-HCl (pH 7.5), 5 mM MgCl₂, 4 mM ATP, 0.1% NP-40, and 1 mM DTT. The lysates were incubated at 4 °C for 30 min with gentle agitation and clarified by centrifugation at 15,000 × g for 10 min at 4 °C. Equal amounts of protein were mixed with native sample buffer (final concentrations: 50 mM Tris-HCl (pH 7.5), 10% (v/v) glycerol and 0.006% xylene cyanol) and subjected to electrophoresis on 4% native polyacrylamide gels at 4 °C. Electrophoresis was carried out at 100 V for 3 h. After electrophoresis, proteins were transferred to PVDF membrane by electroblotting. Membrane were incubated with primary and secondary antibodies using standard procedures, and protein bands were visualized using a NOVEX ECL Chemiluminescent Substrate Reagent Kit (Thermo Fisher Scientific) and an ImageQuant LAS 4000 mini (GE Healthcare).

### Measurement of mitochondrial membrane potential

Mitochondrial membrane potential was evaluated using the MT-1 MitoMP Detection Kit (Dojindo Laboratories, Kumamoto, Japan) or MitoTracker Red CMXRos (Thermo Fisher Scientific, Waltham, MA, USA) according to the manufacturer’s instructions. HeLa cells seeded on glass-bottom dishes were incubated with MT-1 or MitoTracker Red diluted 1:1000 in 1× working solution provided in the kit for 30 min at 37 °C. After washing with 1× working solution, the dish filled with same solution was then mounted on an SP8 confocal microscope (Leica, Wetzlar, Germany) maintained at 37 °C with 5% CO_2_ (BLAST, Kanagawa, Japan). Images were acquired every minute for a total of 7 min, and fluorescence intensity was quantified using LAS X software (Leica).

### Morphological analysis of mitochondria

HeLa cells were co-transfected with plasmids encoding mCherry-Sec61B and EGFP-MAO, in which EGFP is fused to the mitochondrial targeting sequence of monoamine oxidase (MAO) ^59^, using FuGENE HD (Promega, Madison, WI, USA) according to the manufacturer’s instructions. One day after transfection, cells were treated with CGK733 and fixed with 4% paraformaldehyde for 10 min. Fluorescence images were acquired using an SP8 confocal microscope equipped with a 63×/1.40 HC PL APO Oil CS2 oil-immersion objective (Leica).

### Statistical analysis

Statistical analyses were performed using R Commander. One-way ANOVA followed by Dunnett’s test was applied to determine statistical significance. Data are presented as mean ± standard deviation (S.D.). The sample size (n) for each experiment is indicated in the figure legends. Differences with *p* < 0.05 were considered statistically significant.

---

**Extended Data Figure 1.**
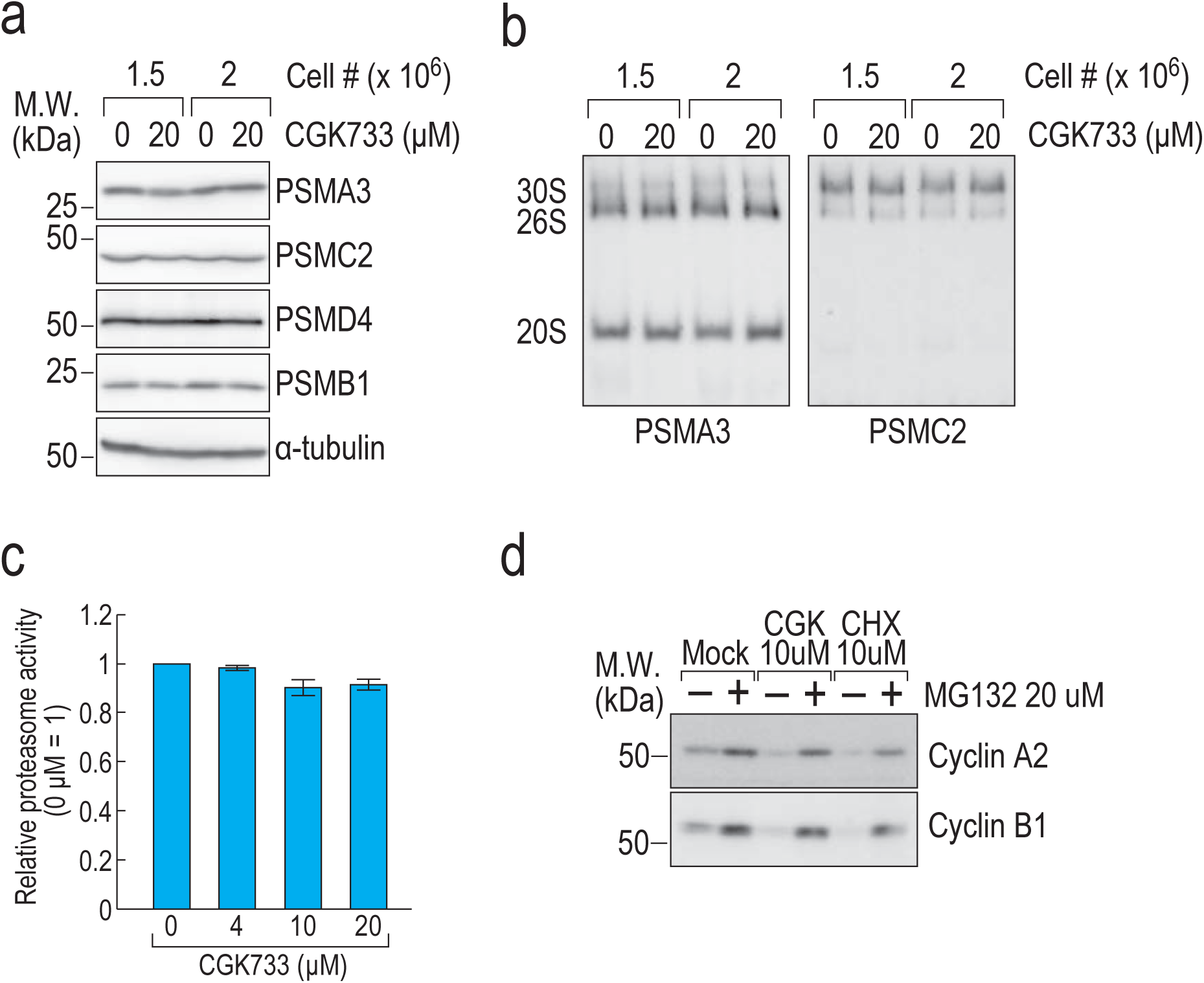
CGK733 does not affect proteasome activity. (a, b) HeLa cells were treated with the indicated concentrations of CGK733 for 6 h. Protein levels were analyzed by immunoblotting with the indicated antibodies (a), and proteasome assembly was assessed by native PAGE followed by immunoblotting with the indicated antibodies (b). (c) Proteasome activity was assessed by in vitro proteasome activity assay (n = 3). Error bars represent s.d. (d) HeLa cells were treated with 10 μM CGK733, 10 μg/ml cycloheximide (CHX), and/or 20 μM MG132 for 6 h. Protein levels were analyzed by immunoblotting with the indicated antibodies.

**Extended Data Figure 2.**
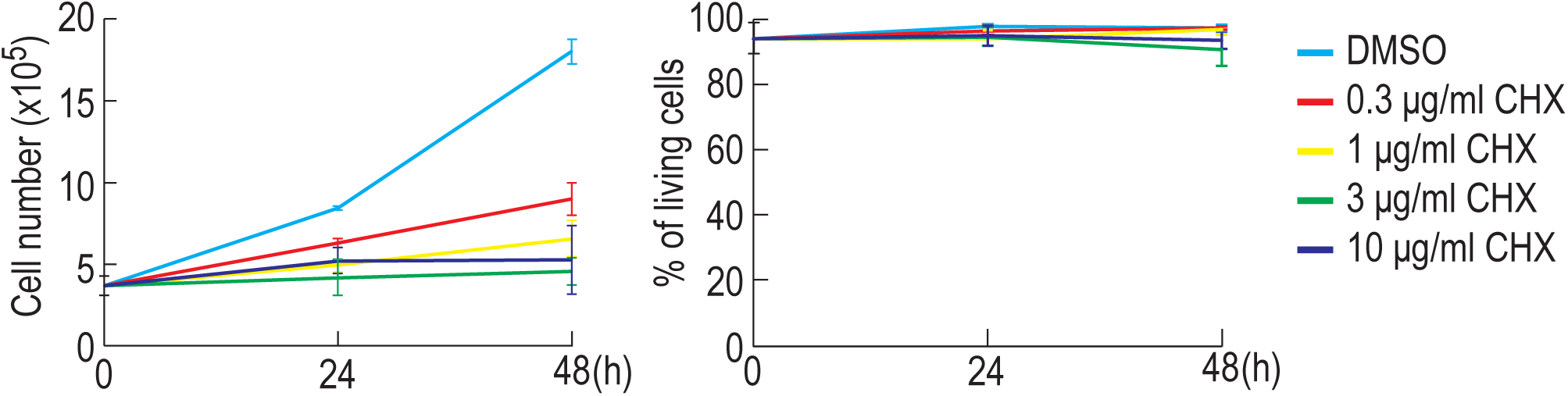
Cycloheximide inhibits cell proliferation but does not induce cell death. HeLa cells were treated with the indicated concentrations of cycloheximide for 24 or 48 h. Cell count and viability were assessed at each time point (n = 3). Error bars represent s.d.

**Extended Data Figure 3.**
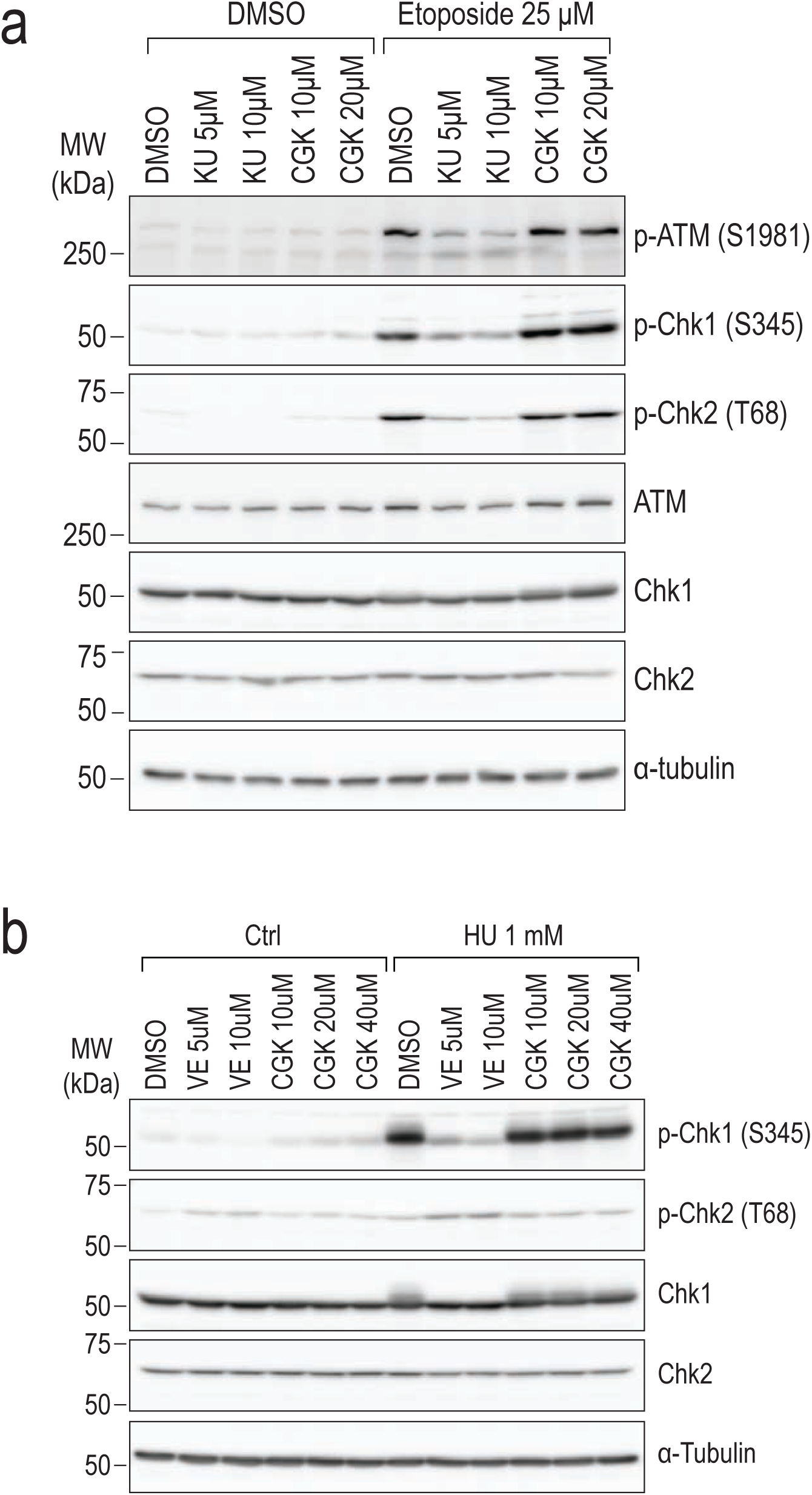
CGK733 does not inhibit ATM/ATR. HeLa cells were treated with the indicated concentrations of hydroxyurea (HU), KU55933, CGK733, etoposide, and/or VE-821 for 6 h. Protein levels were analyzed by immunoblotting with the indicated antibodies.

**Extended Data Figure 4.**
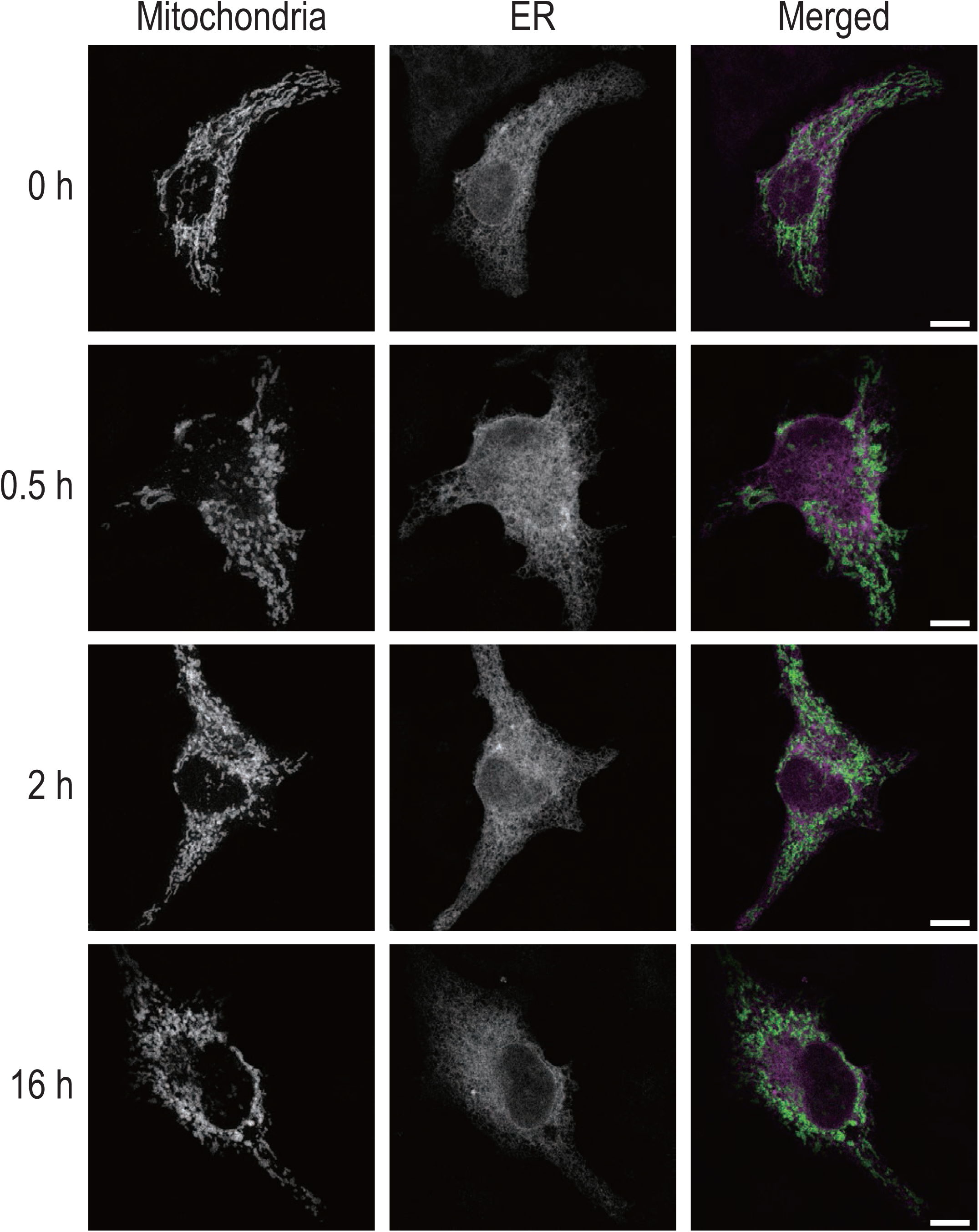
CGK733 induces mitochondrial fragmentation. HeLa cells expressing ER (mCherry-Sec61B) and mitochondrial (EGFP-MAO) markers were treated with 5 μM CGK733 for the indicated times, and the morphology of the ER and mitochondria was examined. Scale bar, 10 μm

**Extended Data Figure 5.**
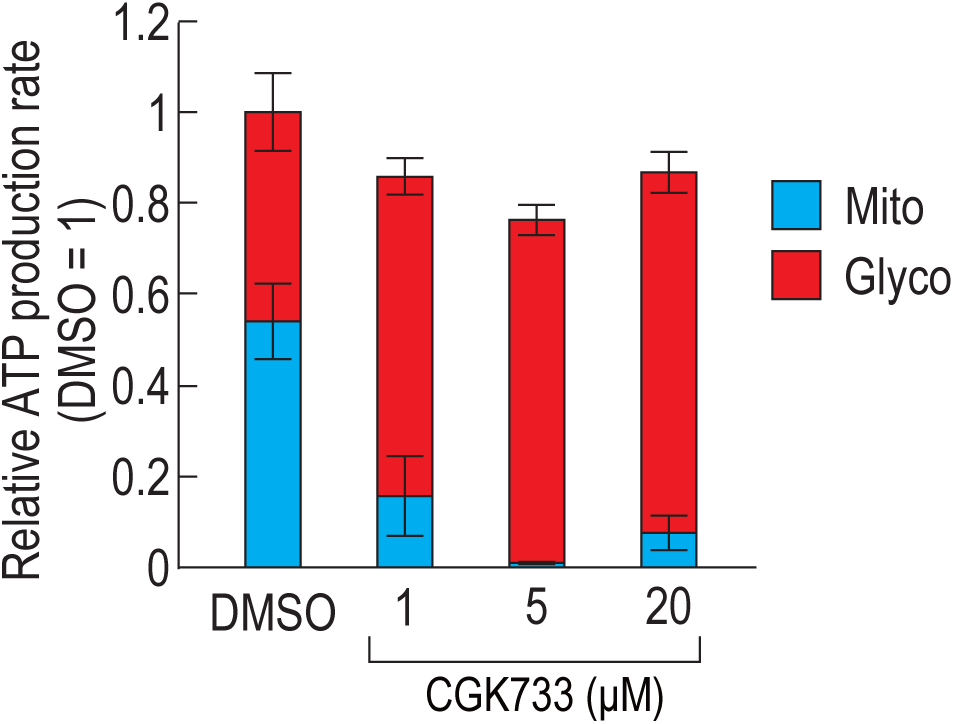
CGK733 induces a metabolic shift from mitochondrial respiration to glycolysis for ATP production. The ATP production rate of CGK733-treated cells was calculated from the data shown in Fig. 5a (n = 3). Error bars represent s.d.

**Extended Data Figure 6.**
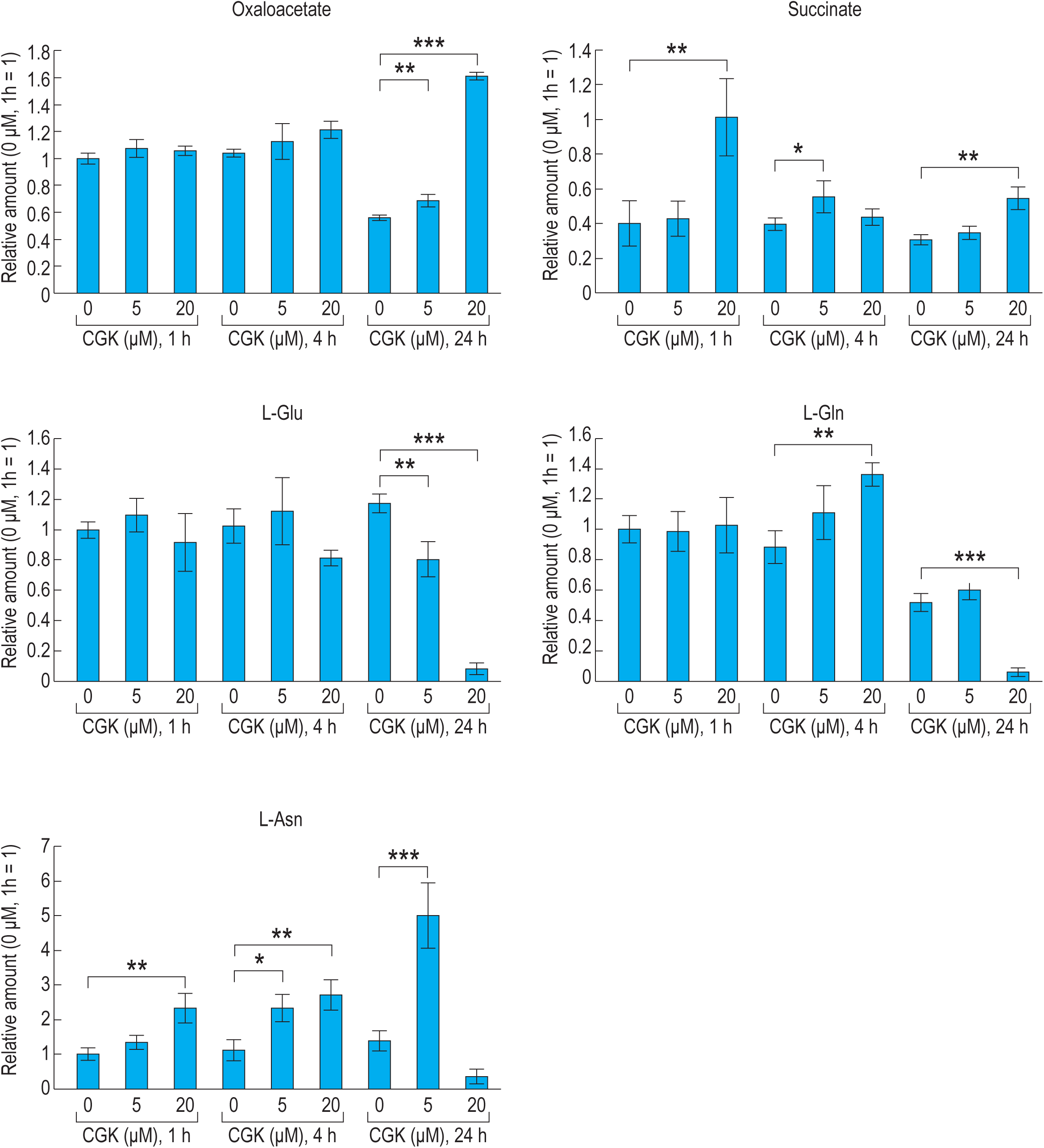
Amount of TCA cycle metabolites and amino acids in CGK733-treated cells. Relative amounts of TCA cycle metabolites and amino acids in cells treated with the indicated concentrations of CGK733 for 1, 4, or 24 h. Statistical significance was determined by one-way ANOVA followed by Dunnett’s test (*: p<0.05; **: p<0.01; ***: p<0.001). Error bars represent s.d.

**Extended Data Figure 7.**
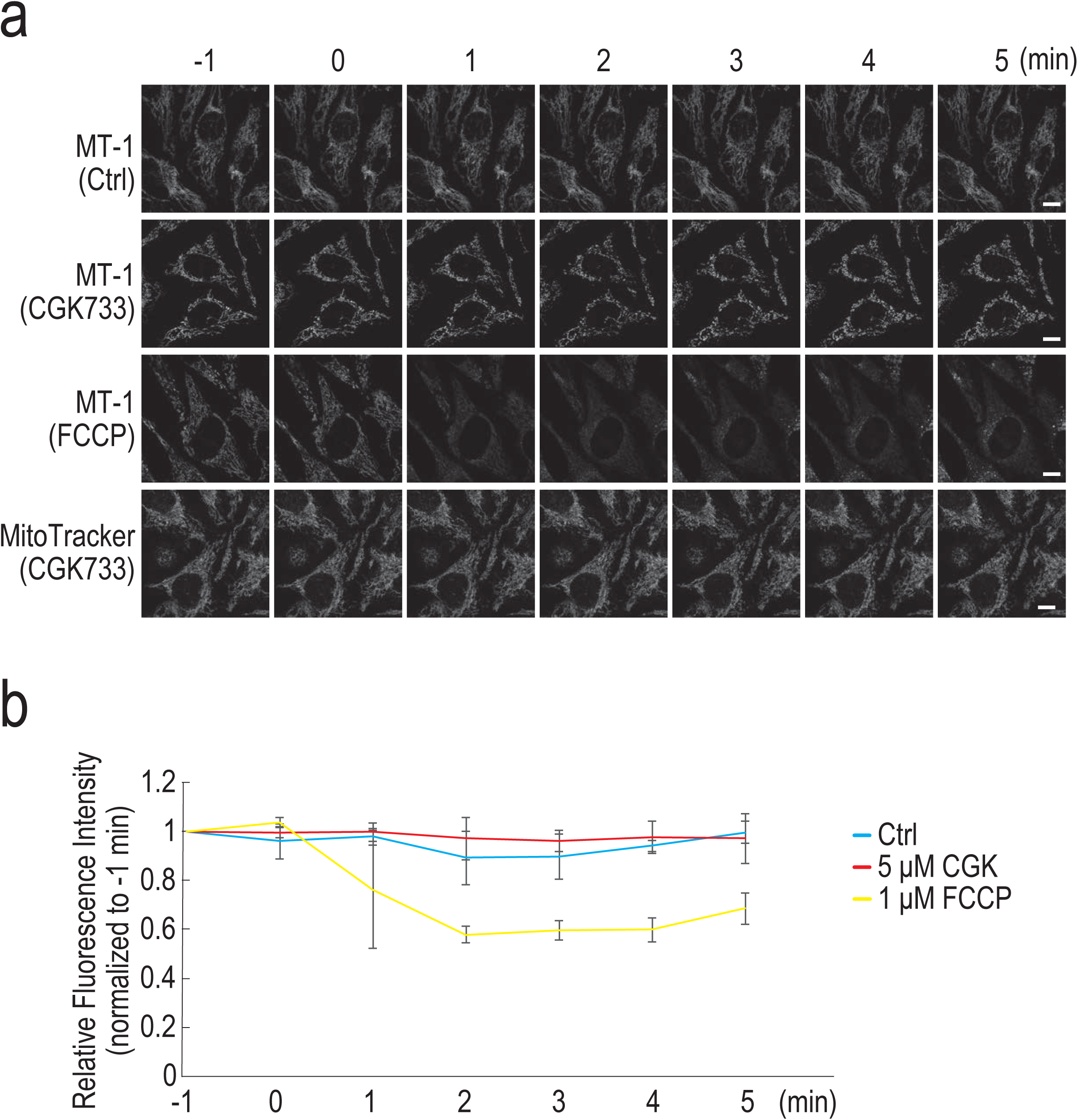
CGK733 does not affect mitochondrial membrane potential. (a) The mitochondrial membrane potential in live cells treated with 5 μM CGK733 or 1 μM FCCP was examined using MT-1 dye or MitoTracker Red dye by time-lapse imaging. CGK733 and FCCP were added at 0 min, and images were acquired every minute. Scale Bar, 10 μM. (b) Relative MT-1 fluorescence intensity was calculated from the data shown in (a) (Ctrl: n = 5; CGK733: n = 6; FCCP: n = 5). Error bars represent s.d.

**Extended Data Figure 8.**
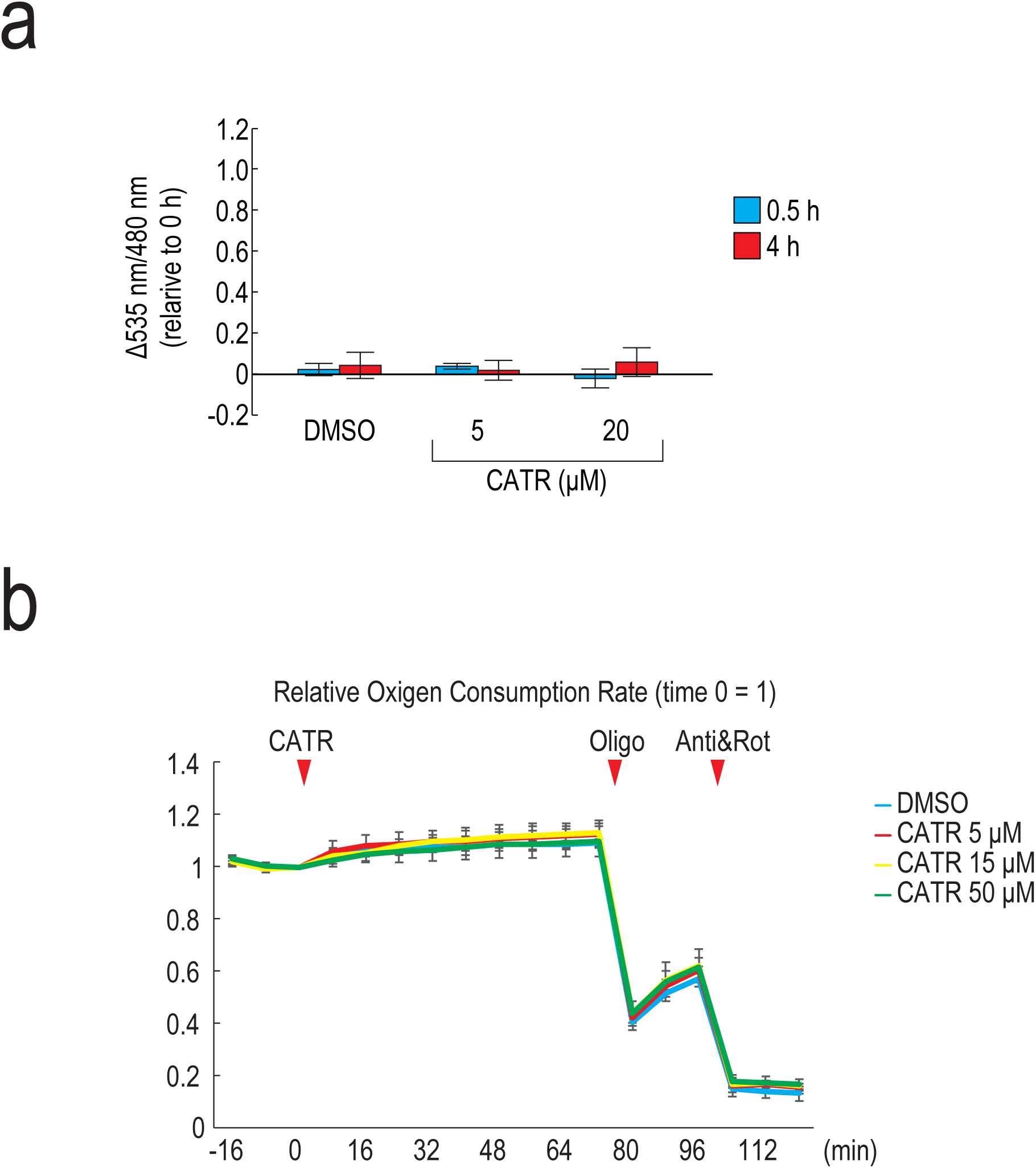
Carboxyatractyloside does not induce ATP accumulation in the mitochondria or affect OCR. (a) HeLa cells expressing mitATeam were treated with 5 µM (n = 11 at 0.5 and 4 h) or 20 µM (n = 10 at 0.5 and 4 h) CATR, or with 0.1% DMSO (n = 21 at 0.5 h; n = 22 at 4 h) for 0.5 or 4 h. Changes in mitochondrial emission ratios relative to 0 h were calculated. (b) OCR of cells was measured after sequential treatment with the indicated concentrations of CATR, 1.5 μM oligomycin (Oligo), and 500 nM rotenone and antimycin A (Rot/Anti). Error bars represent s.d.

**Supplementary Table 1.**
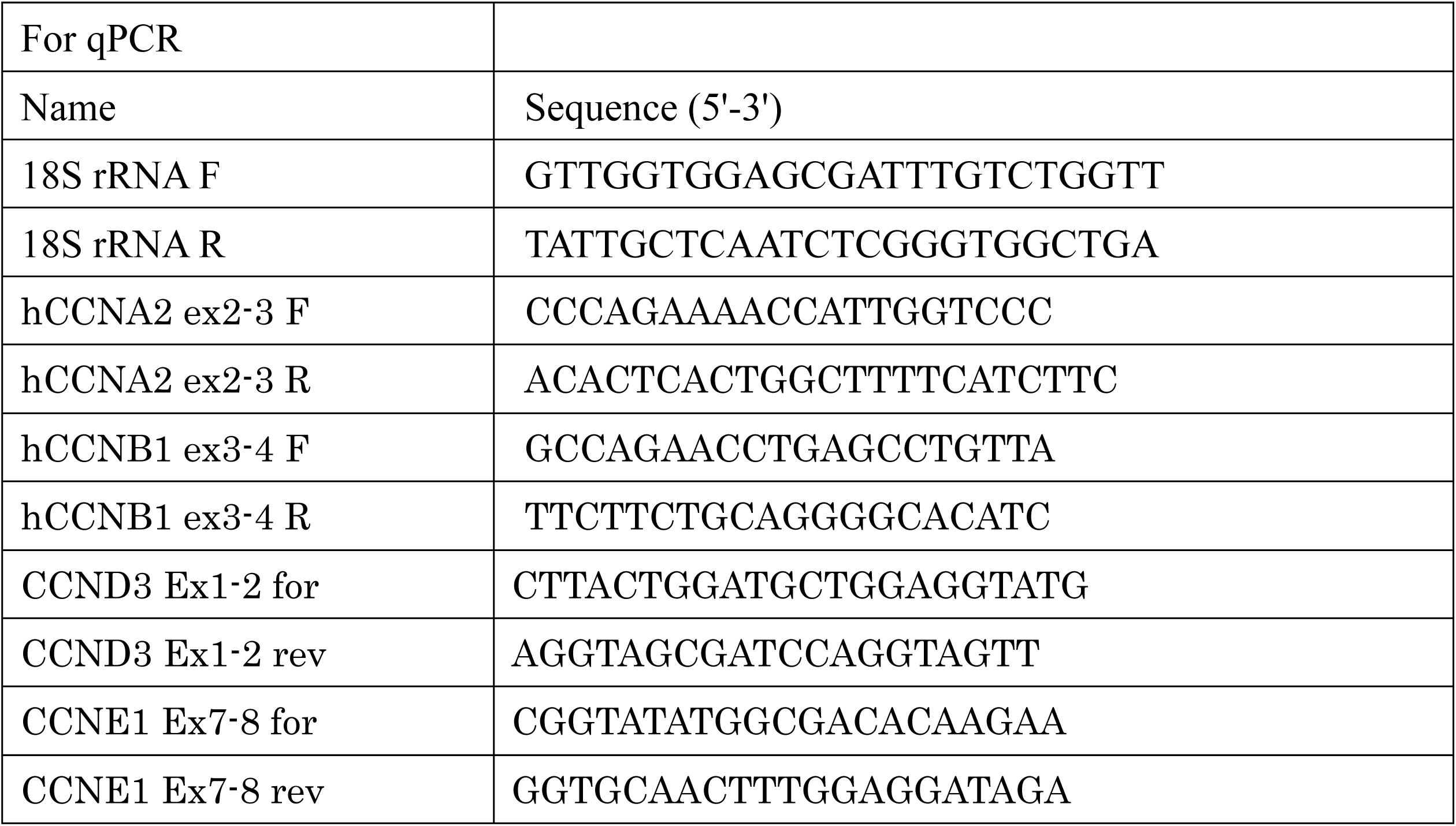
List of primer used in this study.

**Extended Data Table 1.**
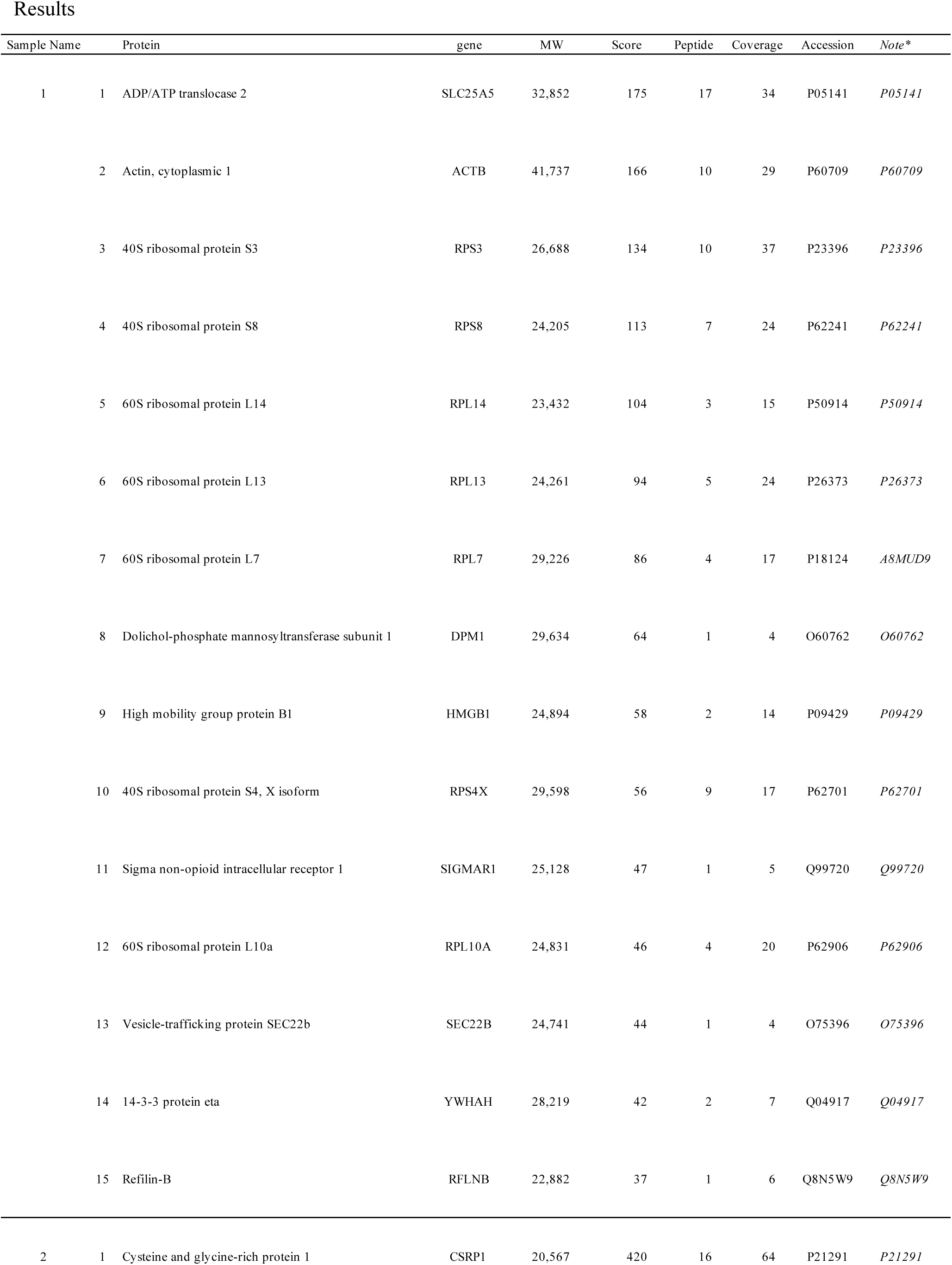
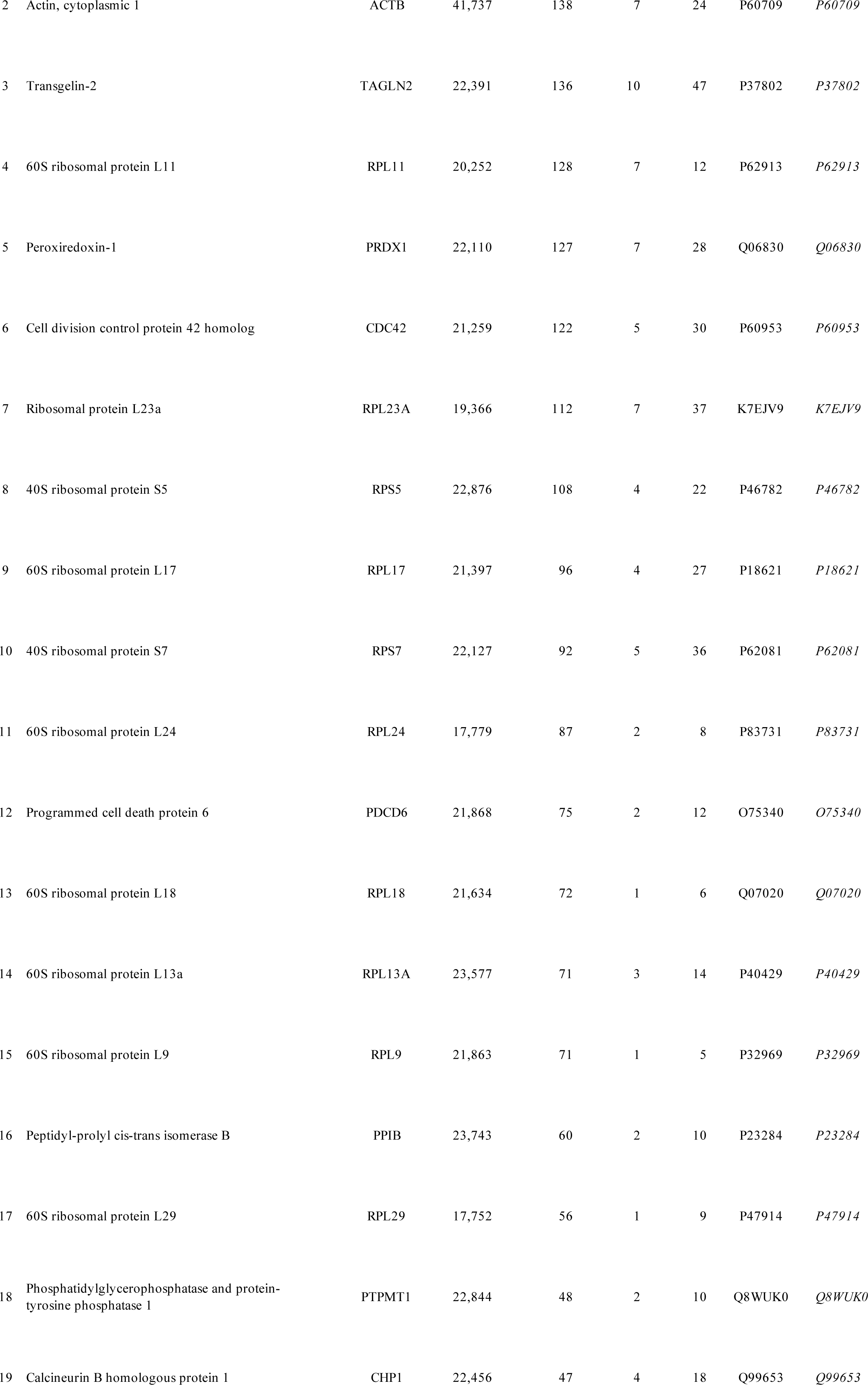
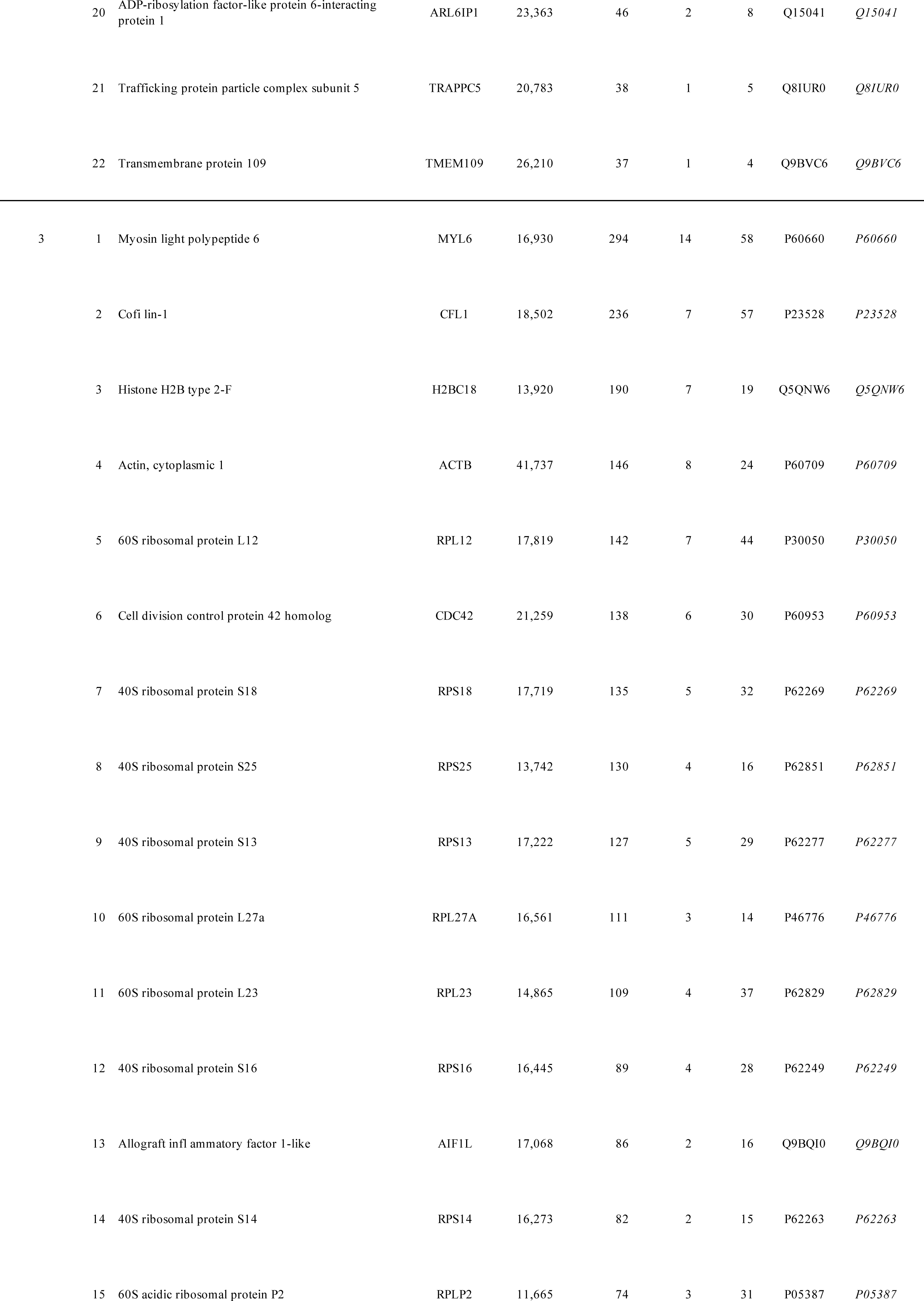
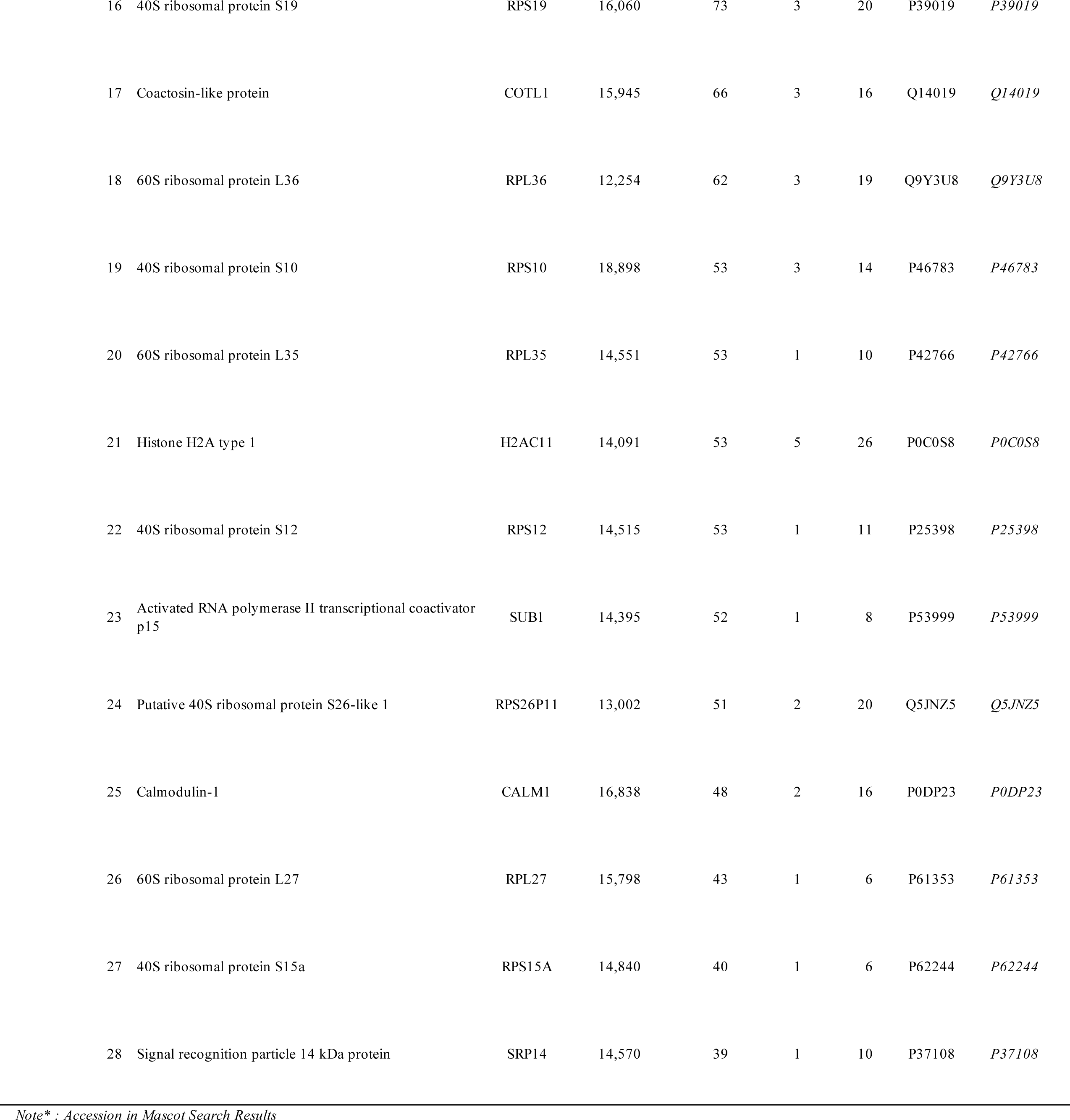
Candidates for CGK733-binding proteins.

